# Temperature impacts the transmission of malaria parasites by *Anopheles gambiae* and *Anopheles stephensi* mosquitoes

**DOI:** 10.1101/2020.07.08.194472

**Authors:** Oswaldo C. Villena, Sadie J. Ryan, Courtney C. Murdock, Leah R. Johnson

## Abstract

Extrinsic environmental factors influence the spatio-temporal dynamics of many organisms, including insects that transmit the pathogens responsible for vector-borne diseases (VBDs). Temperature is an especially important constraint on the fitness of a wide variety of insects, as they are primarily ectotherms. Temperature constrains the distribution of ectotherms and therefore of the infections that they spread in both space and time. More concretely, a mechanistic understanding of how temperature impacts traits of ectotherms to predict the distribution of ectotherms and vector-borne infections is key to predicting the consequences of climate change on transmission of VBDs like malaria. However, the response of transmission to temperature and other drivers is complex, as thermal traits of ectotherms are typically non-linear, and they interact to determine transmission constraints. In this study, we assess and compare the effect of temperature on the transmission of two malaria parasites, *Plasmodium falciparum* and *Plasmodium vivax*, by two malaria vector species, *Anopheles gambiae* and *Anopheles stephensi*. We model the non-linear responses of temperature dependent mosquito and parasite traits (mosquito development rate, bite rate, fecundity, egg to adult survival, vector competence, mortality rate, and parasite development rate) and incorporate these traits into a suitability metric based on a model for the basic reproductive number across temperatures. Our model predicts that the optimum temperature for transmission suitability is similar for the four mosquito-parasite combinations assessed in this study. The main differences are found at the thermal limits. More specifically, we found significant differences in the upper thermal limit between parasites spread by the same mosquito (*An. stephensi*) and between mosquitoes carrying *P. falciparum*. In contrast, at the lower thermal limit the significant differences were primarily between the mosquito species that both carried the same pathogen (e.g., *An. stephensi* and *An. gambiae* both with *P. falciparum*). Using prevalence data from Africa and Asia, we show that the transmission suitability metric *S*(*T*) calculated from our mechanistic model is an important predictor of malaria prevalence. We mapped risk to illustrate the areas in Africa and Asia that are suitable for malaria transmission year-round based temperature.

## 1 Introduction

The physiological processes of many organisms, especially ectotherms, are highly constrained by ambient temperature. In ectotherms, including many insects, body temperatures fluctuate with ambient temperatures. In turn, the rates of most of their biological and biochemical processes shift with temperature, impacting ectotherms fitness (e.g., development rate, survival) (Abram et al., 2017; Kern et al., 2015). Thus temperature contributes to the observed dynamics and distribution of populations of ectotherms.

Predicting impact of temperature on both the dynamics and distribution of these ectotherms now and in the future requires a detailed understanding of the relationship between temperature and performance (Cator et al., 2020). However, experiments can be challenging. Thus much of the best available data are on insects, as they are small, relatively easy to handle, and have short generation times. Further, many insects, such as mosquitoes, are also of public health importance, and as such are well studied. Thus, mosquitoes are a convenient model to investigate the effects of temperature on ectotherms. Further, some of the parasites transmitted by mosquitoes are also ectothermic, allowing the exploration of the impact of temperature on linked systems.

In this paper we explore how temperature impacts the transmission of malaria by *Anopheles* mosquitoes. Malaria, a deadly mosquito-borne disease, is present on five of the world’s seven continents (excluding Australia and Antarctica) (Sinka et al., 2011, 2010, 2012). The World Health Organization reported an estimated 219 million cases of malaria in 91 countries in 2017, an increase of 1.1% over the previous year (World Health Organization, 2018). Of these cases, 435,000 resulted in death. The African region accounts for about 91.5% of malaria cases and mortality, followed by South-East Asian region with 5.1% of cases; the other 3.4% of cases occur in the Eastern Mediterranean Region and the Americas (World Health Organization, 2018). Despite intensive control efforts against malaria for more than a decade, malaria endemicity remains high in much of the world, with high morbidity and mortality, especially in children under 5 years of age (Tizifa et al., 2018).

The *Plasmodium* parasites that cause malaria are spread by Anopheles mosquitoes. Of the 430 Anopheles species, around 30 can transmit malaria. *Anopheles gambiae* is the main malaria vector in Africa (Geissbühler et al., 2007), while *An. stephensi* is an important vector in southern and western Asia (Sinka et al., 2012). Only five of the more than 100 *Plasmodium* species cause malaria in humans: *Plasmodium falciparum, P. vivax, P. malariae, P. ovale*, and *P. knowlesi*. Of these *P. falciparum* and *P. vivax* are the most common of the *Plasmodium* species that cause malaria in humans (Snow et al., 2005). *P. falciparum* is responsible for approximately 92% of the recorded malaria cases worldwide, *P. vivax* is responsible for 4% of malaria cases worldwide, and the other three *Plasmodium parasites* are responsible for the other 4% of malaria cases worldwide (World Health Organization, 2008). 99.7% of the malaria cases in Africa are caused by *P. falciparum*. In contrast, South-East Asia has a combination of types, with ∼63% of cases caused by *P. falciparum* and ∼37% by *P. vivax* (World Health Organization, 2018).

Both *Anopheles* mosquitoes and the malaria parasites they vector are ectothermic organisms (Beck-Johnson et al., 2017; Clements, 2013; Fang and McCutchan, 2002; Lahondére and Lazzari, 2012). Thus the fitness of mosquitoes (e.g., survival, development time), and malaria parasite (e.g., incubation period) are highly temperature sensitive (Paaijmans and Thomas, 2011; Sinclair et al., 2016). Further, we expect interactions between the parasites and their vectors that may also be mediated by temperature. Therefore, when and where malaria will be transmitted will be constrained by environmental temperatures.

In this study, we use a mechanistic model to determine the impact of temperature on a suitability metric, *S*(*T*), that is based on the reproductive number (*R*_0_). We aggregated available data from the literature on the thermal responses of mosquito and parasite traits (e.g., mosquito and parasite development rates) measured across multiple constant temperatures. We fit the thermal response of each component of *S*(*T*) independently to these data using a Bayesian approach (Johnson et al., 2015). We then incorporated the posterior distribution of each component trait into *S*(*T*). We validated these models using data on malaria prevalence in Africa and Asia. The validated models allow us to predict thermal suitability for the transmission of *P. falciparum* and *P. vivax* by *An. stephensi* and *An. gambiae* in space. This is followed by a discussion of how the approach taken compares to other studies and the implications of the results.

## 2 Materials and Methods

### 2.1 Thermal Trait Data

We collected published data from the literature on the thermal responses of the following mosquito traits for nine mosquito species of the *Anopheles* genus that can transmit malaria: bite rate (*a*), mosquito development rate (MDR), proportion of eggs to adulthood (*p*_*EA*_), fecundity in eggs per female per day (EFD), and mosquito mortality rate (*µ*). We also collected published data on the thermal response of the parasite development rate (PDR) for five malaria parasites and on vector competence (*bc*) for all vector/parasite pairs that were available (See SM; Appendix A1). Although malaria is one of the best studied vector-borne diseases, the entire suite of temperature dependent mosquito, parasite, and interaction traits for the mosquito-parasite systems is only available for *An. stephensi* with *P. falciparum*. In other cases data are nearly complete. For example, *An. stephensi* with *P. vivax* is missing only vector competence (see SM; Appendix A1). Others have moderate gaps. *An. gambaie* is missing data for bite rate, parasite development rate with *P. vivax*, and vector competence (with either *P. falciparum* or *P. vivax*). For other mosquito species more than two thermal traits were absent. Thus we focus our analysis on *An. stephensi* and *An. gambiae* as these are the most complete (See SM; Appendix A1). Since even these sets have data gaps in thermal traits, where there is a gap we use traits data available from the closest related species based on similar biologic and ecological characteristics (see SM; Appendix A2).

### 2.2 Modeling the temperature dependence of suitability

Mathematical models of disease systems often use *R*_0_, the basic reproductive number, as a measure of disease transmissibility (Holme and Masuda, 2015). This basic reproductive number gives the average number of secondary cases that one infected individual generates during an infectious period in a susceptible population. The most common parameterizations of *R*_0_ for vector-borne infections are based on the Ross-MacDonald model of malaria transmission (Dietz, 1993). Here we specifically use a formulation that incorporates multiple mosquito traits to approximate the mosquito population size (Johnson et al., 2015; Mordecai et al., 2019, 2017, 2013), that is we assume *R*_0_ is given by:

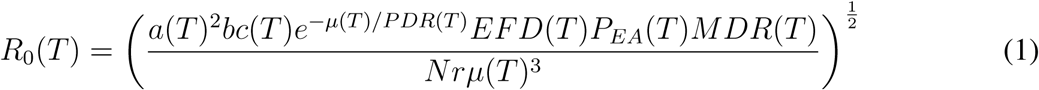

where *a* is the mosquito biting rate; *bc* is vector competence; *µ* is the mosquito mortality rate; *PDR* is the parasite development rate; *EFD* is the mosquito fecundity expressed as the number of eggs per female per day; *P*_*EA*_ is the proportion of eggs to adulthood; *MDR* is the mosquito development rate; *N* is the density of humans or hosts; and *r* is the human recovery rate. Because we are interested in the shape of the thermal response only, we define a suitability metric, *S*(*T*), that only incorporates the temperature dependent components, that is:

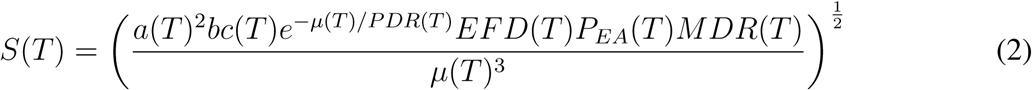

Most thermal traits of ectotherms exhibit unimodal responses (Colinet et al., 2015; Mordecai et al., 2019). Thus we assume that all of the components of *S*(*T*) are temperature dependent. For each of these individual traits (e.g., bite rate) for each mosquito species we fit one of three kinds of unimodal thermal response. For asymmetric responses like *MDR, a*, and *PDR* we fit a Briére function:

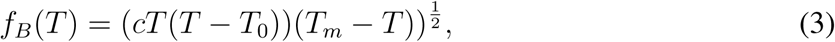

where *T*_0_ is the lower thermal limit (where the response becomes zero), *T*_*m*_ is the upper thermal limit, and *c* is a constant that determines the curvature at the optimum. Formally we assume a piecewise continuous function, so that the thermal trait is assumed to be zero if *T* < *T*_0_ or *T* > *T*_*m*_. Symmetric responses come in two flavors: concave down or concave up. For concave-down symmetric responses like *bc, P*_*EA*_, and *EFD*, we fit a quadratic function parameterized in terms of the temperature intercepts,

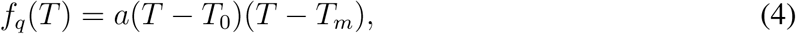

where *T*_0_ is the lower thermal limit, *T*_*m*_ is the upper thermal limit, and *a* is a constant that determines the curvature at the optimum. As with the Briére function we assume the trait is piecewise zero above and below the thermal limits. For concave-up symmetric responses like *µ*, we fit a concave-up quadratic function

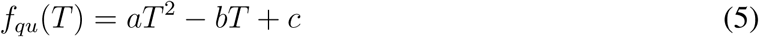

where *a, b*, and *c* are the standard quadratic parameters. Note that because all traits must be ≥ 0 we also truncate this function, creating a piecewise continuous function where *f*_*qu*_ is set to zero if the quadratic evaluates to a negative value.

#### 2.2.1 Bayesian Fitting of Thermal Traits

We fit each unimodal thermal response for all traits for each mosquito species (*An. stephensi* and *An. gambiae*) or parasite strain (*P. falciparum* and *P. vivax*) with a Bayesian approach using the JAGS/`rjags` package (Plummer, 2016) in R (R Development Core Team, 2017). We defined an appropriate likelihood for each trait (e.g., binomial likelihoods for proportion data, truncated normal for continuous numeric traits) with the mean defined by the either a Briére function (for asymmetric relationships) or quadratic (symmetric relationships). For all traits we chose relatively uninformative priors that limit each parameter to its biologically realistic range. More specifically, we assumed that temperatures below 0°C and above 45°C are lethal for both mosquitoes and malaria parasites (Lyons et al., 2012; Mordecai et al., 2017). Based on these assumptions we set uniform priors for the minimum temperature (*T*_0_) between 0 – 24°C and for the the maximum temperature (*T*_*m*_) between 25 – 45°C. Priors for other parameters in the thermal responses were set, to ensure parameters were positive and not tightly constrained (See SM; Appendix A3). The `rjags` package uses a Metropolis within Gibbs Markov Chain Monte Carlo (MCMC) sampling scheme to obtain samples from the joint posterior distribution of parameters. For each fitted trait, we obtained posterior samples from five Markov chains that were run for 20000 iterations initiated with random starting values. These samples were obtained after using 10000 iterations for adaptation and burning another 10000 iterations. We visually assessed convergence of the Markov chains. To obtain the posterior summaries of each trait, we combined the 20000 samples from each Markov chain from the posterior distribution, which resulted in a total of 100000 posterior samples of the thermal response for each trait. Based on these samples, for each unimodal thermal response we calculated the posterior mean and 95% highest probability density (HPD) interval around the mean of the thermal response and various summaries (e.g., the thermal minimum or maximum).

Once the posterior samples of parameters for all thermal traits across temperature for each species were obtained, these curves were combined to produce 100000 posterior samples of *S*(*T*) (See SM; Appendix A4 for the traits used for each mosquito/parasite set). We used these samples of *S*(*T*) across temperature to calculate the posterior mean and the 95% HPD of the overall thermal response of suitability, as well as for the critical thermal minimum, thermal maximum, and optimal temperature for transmission suitability by the two mosquito species for each parasite. We also used the posterior samples of the extrema of *S*(*T*) (lower thermal limit, upper thermal limit, and optimum) to assess the significance of the differences in these summaries between the four mosquito/parasite combinations. For all of the possible pairwise mosquito/parasite sets, we calculated the probability that the differences between each pair of samples of the *S*(*T*) posterior distributions for the mosquito-parasite combinations is greater than zero. The algorithm to compute this based on posterior samples is as follows. 10000 random samples were selected from the posterior distribution of the target statistics of the *S*(*T*) (e.g., the thermal minimum) from each mosquito/parasite combination. One mosquito/parasite pair is chosen as the baseline, and a second as the comparator. The difference between each sample of the baseline mosquito/parasite pair and the comparator are calculated, resulting in 10000 differences. If the two sets of samples were random draws from the sample distribution, we would expect to see approximately even numbers of differences above and below zero, and the mean would be close to zero. We calculated the mean of the differences (primarily to identify which of the baseline or comparative was on average higher, with a positive value indicating that the baseline was larger). We then calculated the proportion of differences that skewed towards the baseline. If the proportion of differences in the direction of the baseline is greater than or equal 0.95, we classify this as a significant difference in the statistic or there is strong evidence that the values are different. If the proportion of that the differences skewed in the direction of the baseline is between 0.6 and 0.94, we said that there is some evidence that the difference is significant. Otherwise we said that there is not a significant difference, because the difference between these two groups could be explained by chance alone.

We also performed an uncertainty analysis for *S*(*T*) to quantify the contribution of each trait to the overall uncertainty in suitability. We calculated the uncertainty due to each thermal trait on *S*(*T*) as follows. First we set all traits but one focal trait to their respective posterior means. We then recalculated the conditional posterior distribution of *S*(*T*) with only the focal trait allowed to vary according to its full posterior distribution. Then at each temperature we calculated the width of the 95% HPD due to only the variation in this single parameter. We calculated the relative width of the HPD interval for this focal trait compared to the HPD interval when all traits are allowed to vary according to their full posterior distribution. This is approximately the proportion of the uncertainty each parameter contributes to the full uncertainty in *S*(*T*) (Johnson et al., 2015). This process was repeated for each focal trait in turn.

### 2.3 Model validation

To validate our models, we tested if *S*(*T*) is consistent with data on observed malaria prevalence in Africa and Asia, using generalized linear models (GLMs). We obtained data on *P. falciparum* and *P. vivax* prevalence data collected at the village level in 46 countries in Africa and 21 countries in Asia from 1990 to 2017 from the Malaria Atlas Project (Pfeffer et al., 2018). The prevalence data were matched with monthly aggregated mean temperature data for the quarter prior to the start month of each study obtained from the WorldClim Global Climate Data Project (Fick and Hijmans, 2017) using latitude and longitude coordinates as merging points.

We also included two socio-economic predictor variables in our analysis: adjusted *per capita* gross domestic product (GDP) and population density (*p*). These allow us to account for other potential sources of country to country or local variation in prevalence. Approximate population density data at the village level where the prevalence studies took place were obtained from the Global Rural-Urban Mapping Project - GRUMP project (Balk et al., 2006). GRUMP raster data containing population density data were imported into ArcGIS software version 10.8 and population density data at each location in our prevalence dataset was extracted using the “extract values to points” tool from the spatial analysis toolset in ArcMap (Scott and Janikas, 2010). GDP data were obtained from the Institute for Health Metrics and Evaluation of the USA (James et al., 2012). Both population density and GDP across countries exhibit clumpiness and variation across orders of magnitude, which can lead to high leverage data points having outsized influence on results. Logarithmic transformation of variables is a very common practice for addressing these issues with predictors (Changyong et al., 2014). Thus, for all of our analyses we used the natural logarithm transformed versions of both of population density and GDP.

We categorized each location in our dataset with a binary response variable to indicate whether or not malaria had been observed at that location in our prevalence dataset, that is we defined the response *y*_*i*_ for the *i*^th^ location as

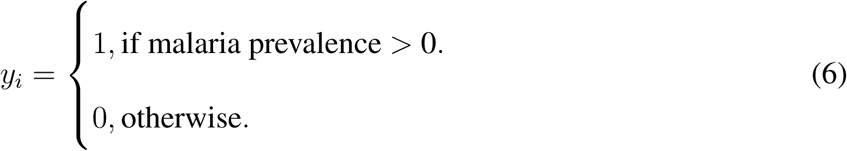

We fit a series of logistic regression models with different subsets of predictors that include *S*(*T*), log population density, and log GDP, or a combination of these (See SM; Appendices A9 and A10 for full models). The general logistic GLM is defined by the mean equation:

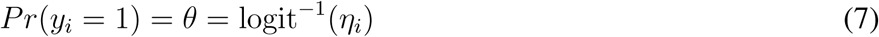

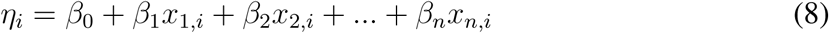

where *η*_*i*_ is the linear predictor, with *β*_0_ as the intercept, *β*_1_, …, *β*_*n*_ are the regression parameters, and *x*_1_, …, *x*_*n*_ are the explanatory variables. Observations at data point *i* are then Binomial random variables with “success” probability *θ* and sample size *n*_*i*_ at each location as defined in Eq. 7. In addition to log *p*, log GDP, and *S*(*T*) we define additional combinations and functions of these parameters to use as potential predictors. First we define *S*(*T*)GTZ, which is the posterior probability that *S*(*T*) > 0. We also defined multiplicative combinations, to account for possible interactions, specifically log(S(T)*GTZ* × GDP) and log(S(T)*GTZ* × *p*). In our models we also consider more standard interactions between log(*p*), log(GDP) and *S*(*T*) > 0. To assess if model assumptions were adequately met we computed and plotted the randomized quantile residuals (RQRs), implemented in the `statmod` package (Giner and Smyth, 2016). RQRs are based on the idea of inverting the estimated distribution function for each observation to obtain standard normal residuals if model assumptions are met and the model fits adequately. RQRs are the residuals of choice for GLM models in large dispersion situations (Dunn and Smyth, 1996). We then chose between all candidate models in terms of model probabilities (*Pr*) that are based on the Bayesian information criterion (BIC) values. Given a list of possible models (*M*_1_, *M*_2_, …, *M*_*i*_), the probability that a model *i* is the best is giving by the equation:

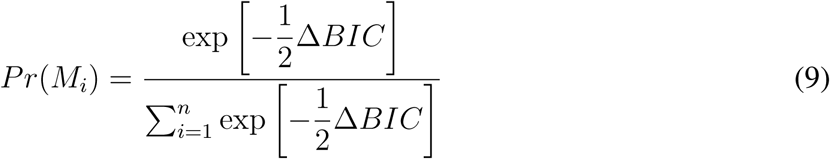

where Δ*BIC* = *BIC*_(*M*)_ − min (*BIC*_(*M*)_)) is the difference between the BIC value for the i^*th*^ model, *BIC*_*Mi*_, and the value of the lowest BIC in the calculated set, min(*BIC*_(*M*_*i*)). The best model will be the one with the highest model probability (Burnham and Anderson, 2004). We also calculated the proportion of deviance explained from each model (*D*^2^). A perfect model has no residual deviance and its (*D*^2^) will be equal to 1. The (*D*^2^) is the GLM analogue to (*R*^2^) in linear models (Dunn and Smyth, 2018).

Following methodology for spatial validation of suitability prediction models in (Taylor et al., 2019; Tesla et al., 2018), we also calculated the proportion of *P. falciparum* and *P. vivax* confirmed positive prevalence cases for Africa and Asia that falls into the months that are suitable for malaria transmission (0 to 12). The proportion of confirmed positive prevalence cases were calculated in areas that are suitable for either *An. stephensi* or *An. gambiae* in Africa and Asia (See SM; Appendices B19 amd B20).

### 2.4 Mapping temperature suitability for malaria transmission

We chose to focus on illustrating high suitability in Africa and Asia (where malaria is most prevalent). We identified the temperatures defining the top quartile of the posterior median of *S*(*T*) and, at each location in space, we calculated the number of months that the pixel was within these bounds for each mosquito-parasite system. Monthly mean temperature rasters at a 30 second resolution were downloaded from the Worldclim-Global Climate Data project (Fick and Hijmans, 2017) using the `raster` package (Hijmans et al., 2015) in the R environment (R Development Core Team, 2017). We cropped the temperature raster maps to the Africa and Asia continents using the *Crop* function from the `raster` package and assigned values of one and zero depending on whether the probability of *S*(*T*) >0 exceeded the threshold at the temperature in those cells. Maps were created in ArcMap 10.7.1 (Barik et al., 2017).

## 3 Results

### 3.1 Posterior distributions of thermal traits

In Figures 1 and 2, we show the posterior mean and the 95% HPD interval around the mean for *An. stephensi* mosquito and parasite thermal traits (See SM; Appendices B1 and B2 for *An. gambiae*). This enables us to visualize the extent of uncertainty around the mean thermal response. In general, we notice that the uncertainty was greater in the thermal responses for traits of *An. gambiae* mosquitoes than *An. stephensi* mosquitoes. This is largely due to fewer data available for estimating the responses of traits to temperature for this species. More specifically, for *An. gambiae*, mosquito development rate (MDR), number of eggs per female per day (EFD), parasite development rate with *P. falciparum* (PDR), and vector competence with *P. falciparum* (bc) showed the greatest uncertainty, primarily due to lack of data (See SM, Appendices B1 and B2). Uncertainty in MDR, EFD, and *bc* have considerable uncertainty in the lower end of the thermal response because there are few or no data available below approximately 20°C. PDR has few overall data points, resulting in significant overall uncertainty, but lack of higher temperature data leads to especially large uncertainty in the upper thermal curve/limit. Bite rate (*a*) similarly, has more uncertainty at the upper thermal limit, although the effect is less pronounced because more total data are available.

**Figure 1:**
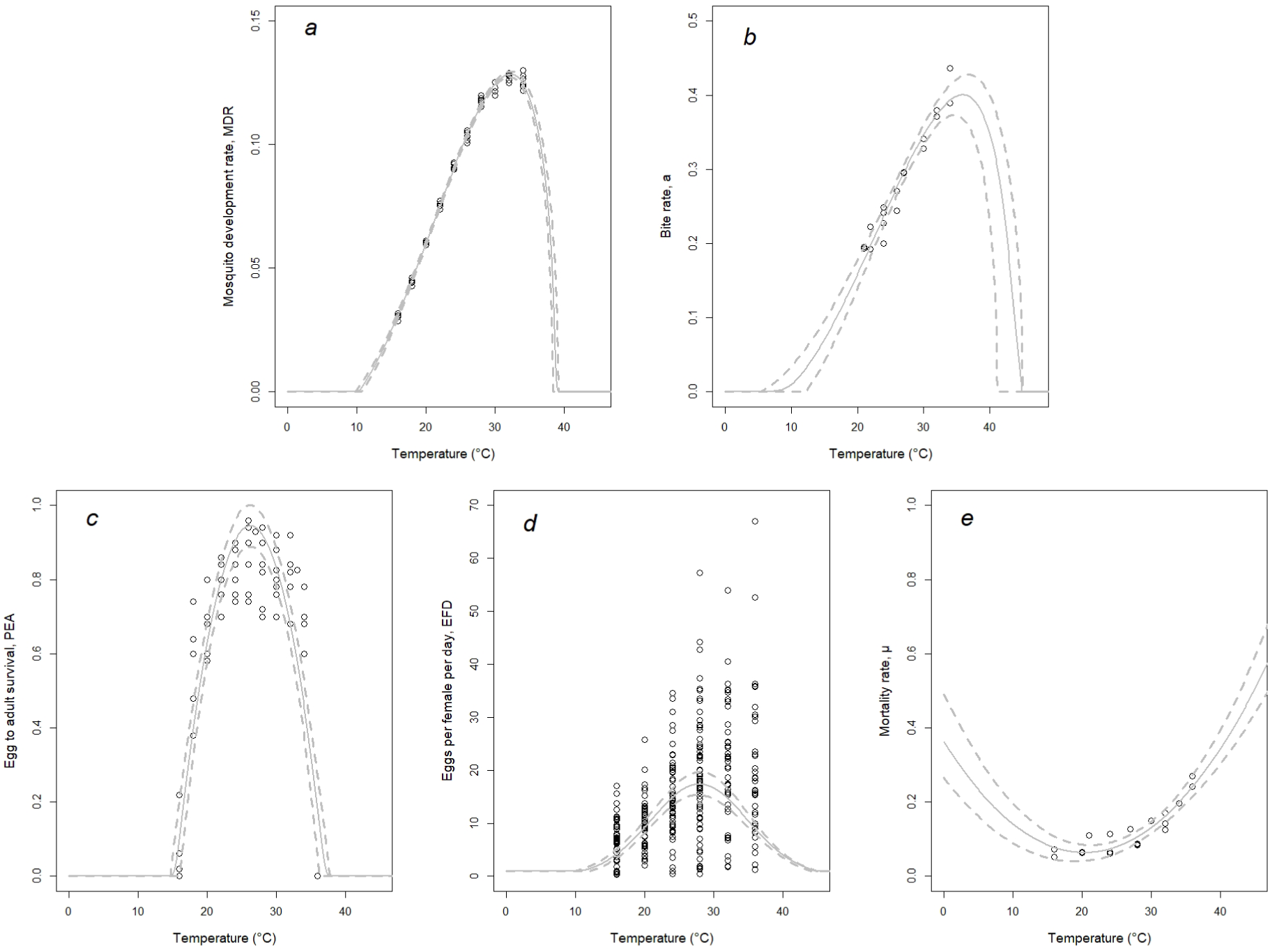
Posterior mean (solid line) and 95% highest posterior density-HPD (dashed lines) of the thermal responses for mosquito traits to calculate *S*(*T*) for *Anopheles stephensi*/ *Plasmodium falciparum* and *An. stephensi*/*P. vivax*. Traits modeled with a Briére thermal response are (A) mosquito development rate and (B) bite rate. Traits modeled with a concave down quadratic function are (C) proportion of eggs to adult and (D) fecundity; and (E) mortality rate which is modeled with a concave-up quadratic function. Data symbols correspond to the species of mosquitoes. ○ is for *An. stephensi*.

**Figure 2:**
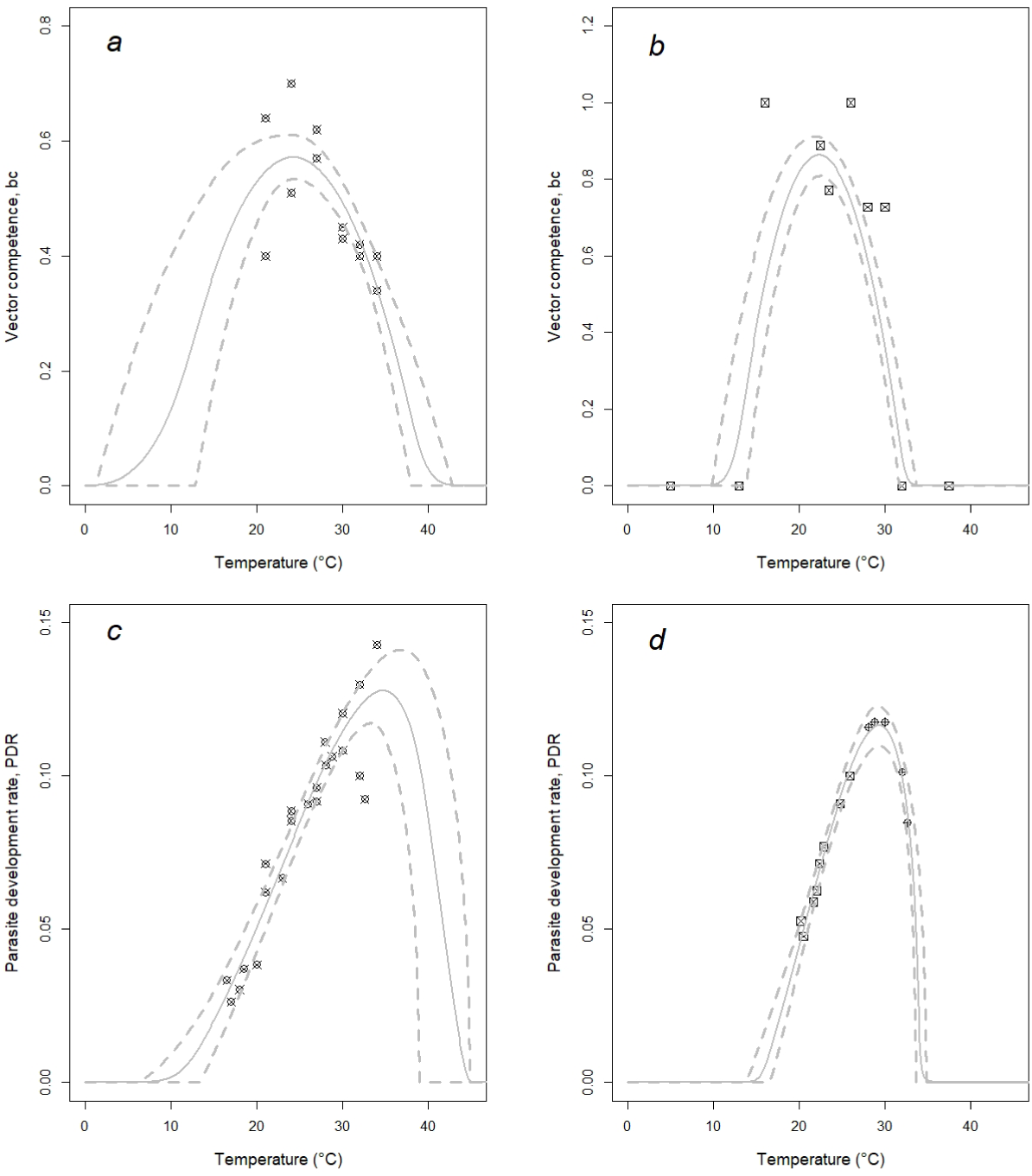
Posterior mean (solid line) and 95% highest posterior density-HPD (dashed lines) of the thermal responses for mosquito/parasite traits to calculate *S*(*T*) for *Anopheles stephensi*/*Plasmodium falciparum* and *An. stephensi*/*P. vivax*. Traits modeled with a concave down quadratic function are vector competencen(A) *P. falciparum* (B) *P. vivax*. Traits modeled with a Briére thermal response are parasite development rates (C) *P. falciparum* (D) *P. vivax*. Data symbols correspond to the species of mosquitoes and parasite used for the analysis. ⊗ is for *An. stephensi* in combination with *P. falciparum*, ⊕ is for *An. stephensi* in combination with *P. vivax*, and ⊠ is for *An. quadrimaculatus* in combination with *P. vivax*

In contrast, most *An. stephensi* traits exhibit considerably less uncertainty around the mean. Where uncertainty exists, for example in the thermal limits of bite rate (*a*), (Figure 1B), and mortality (*µ*), (Figure 1E)), this seems again to be due primarily to a lack of data near the thermal extremes. Similarly, parasite traits with *An. stephensi* mosquitoes typically have better data coverage than *An. gambiae*, and so have less uncertainty. However vector competence, *bc*, for *An. stephensi* with *P. falciparum* is more uncertain at the minimum thermal limit because data were only available above 21°C (Figure 2A). Similarly PDR for *P. falciparum* is uncertain near the maximum thermal limit because there are only three data points above 32 °C (Figure 2C). In contrast *P. vivax* related traits are better constrained.

### 3.2 Posterior distribution of *S*(*T*)

In Figure 3, we show the posterior mean and 95 % HPD of *S*(*T*) for each of the four mosquito-parasite combinations. To allow for more direct comparison, all curves are scaled to the maximum value of the posterior median *S*(*T*) curve, so the maximum value of the median for each individual curve is one. Overall, uncertainty (measured in the width of the HPD intervals after scaling) is greater for *An. gambiae* with *P. falciparum* and *P. vivax* compared to *An. stephensi* with *P. falciparum* and *P. vivax* (Figure 3). Based purely on posterior median values, *An. stephensi* mosquitoes had the greatest temperature range for transmission suitability, with a range of 15.3 °C to 37.2 °C for *P. falciparum* and 15.7 °C to 32.5 °C for *P. vivax*. The median predicted temperature ranges for the suitability transmission of *P. falciparum* and *P. vivax* by *An. gambiae* mosquitoes are very similar with ranges from 19.1 °C to 30.1 °C and 19.2 °C to 31.7 °C respectively (Figure 3). However, these median ranges mask a great deal of uncertainty. The suitability metric for all four mosquito parasite pairs is predicted to peak (i.e., to have optimum) at approximately 25°C. Although there is some variability in this estimate, the posterior distributions of the optimum do not exhibit significant differences between then (See SM; Appendices B13 and B16). In contrast, there are indications that the upper and lower thermal limits may not be the same across the 4 pairs. In *An. stephensi*, there is strong evidence of a difference in the upper thermal transmission limit for these mosquitoes when they transmit *P. falciparum* (CI: 36.5-38 °C) versus *P. vivax* (CI: 31.7-33.7 °C) (See SM; Appendices A7, B12a, and B15a). Similarly, there is strong evidence for differences in the upper thermal limit for transmission of *P. falciparum* when comparing between *An. stephensi* and *An. gambiae* (See SM; Appendices A7, B12c, and B15c). The other comparisons at the maximum thermal limit are not significant (See SM; Appendices B12 and B15).

**Figure 3:**
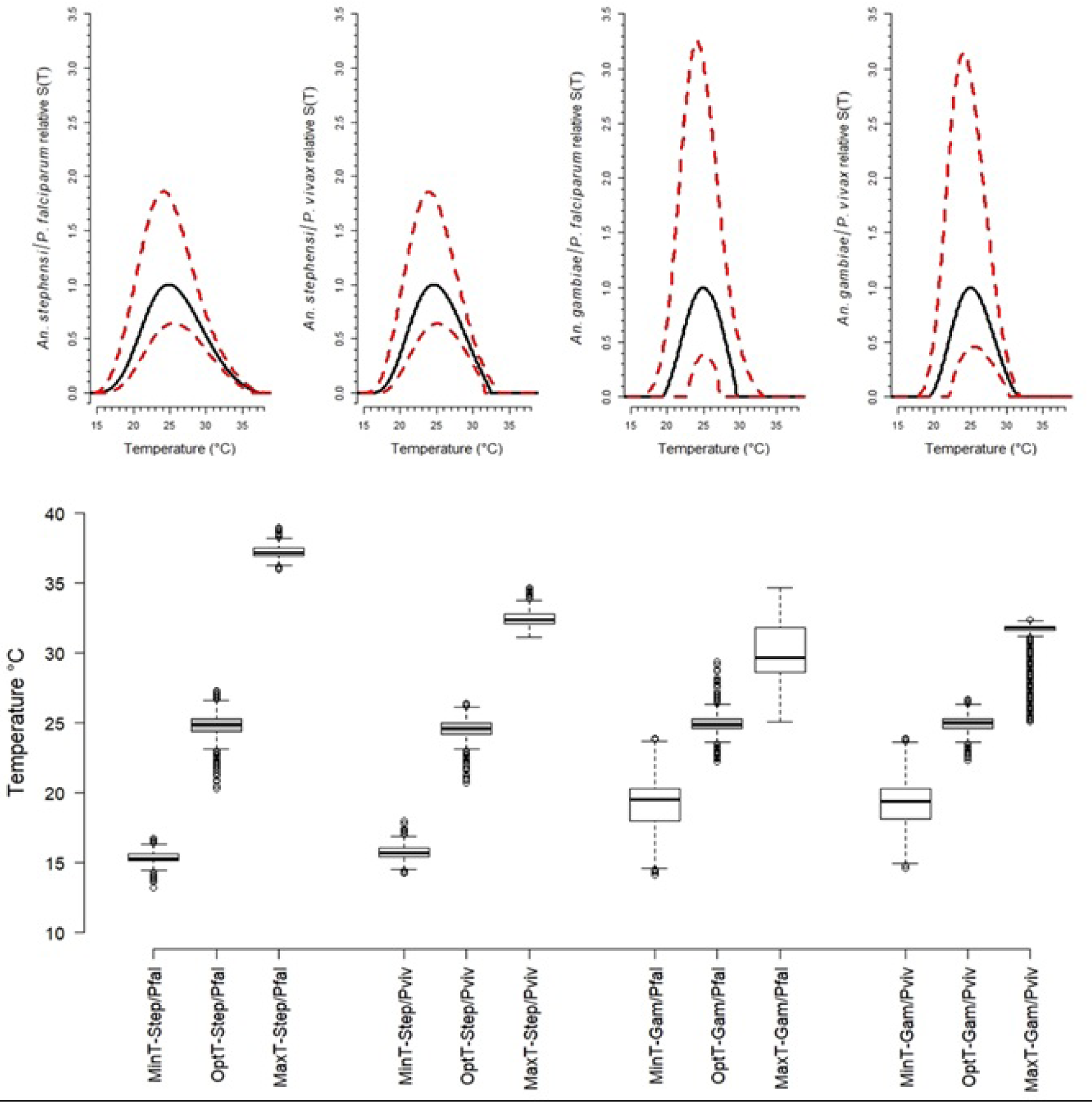
Top row: Relative *S*(*T*) divided by the maximum value of the posterior mean for *An. stephensi* with *P. falciparum, An. stephensi* with *P. vivax, An. gambiae* with *P. falciparum*, and *An. gambiae* with *P. vivax* (from left to rigth). Bottom row: Mean, interquartile range, minimum and maximun numbers from the posterior for the minimum temperature (Min), optimun temperature (Opt), and maximun temperature (Max) for the *R*_0_ of *Anopheles stephensi*/*Plasmodium falciparum* (StepPfal), *An. stephensi*/*P. vivax* (StepPviv), *An. gambiae*/*P. falciparum* (GambPfal), and *An. gambiae*/*P. vivax* (GambPviv).

The predictions for the lower thermal limits were much more similar to each other. Our results show that the only significant differences are between the transmission of *P. falciparum* by *An. stephensi* and by *An. gambiae* mosquitoes (See SM, Appendices B14c and B17c) and between the transmission of *P. vivax* by *An. stephensi* and by *An. gambiae*. The other comparisons at the minimum thermal limit are not significant (See SM; Appendices B14d and B17d).

### 3.3 Sources of uncertainty in *S*(*T*)

Across all combinations of mosquitoes and parasites the uncertainty in *S*(*T*) at intermediate temperatures is dominated by the uncertainty in the adult mosquito mortality rate, *µ* (See SM, Appendices B5 and B6). This is a common pattern as *S* ∝ *µ*^−3^, so small changes in *µ* when *µ* is small (that is near optimal temperatures for mosquito longevity) will have an out-sized impact on *S*. This component is thus almost completely responsible for the location and height of the peak of suitability.

In contrast, the traits that drive uncertainty in the temperatures around the thermal limits varies between each mosquito-parasite pair and is sensitive to the amount and quality of data available for each. For example, our suitability metric for *P. falciparum* in *An. stephensi* seems the most well-resolved of all of the combinations. In this case, uncertainty in *µ* dominates across almost all temperatures, and it is only at the high temperature end, above ≈ 37°C, that most of the uncertainty is caused by something else, specifically egg to adult survival (*P*_*EA*_). At temperatures between 32-37°C vector competence (*bc*) and parasite development rate (PDR) also contribute to the uncertainty, although they do not dominate over either *µ* or *P*_*EA*_. Similarly near the lower temperature regime PDR and *P*_*EA*_ both contribute to the overall uncertainty, but do not dominate compared to *µ* (See SM; Appendix B5c). Because the mosquito traits are shared, the patterns seen in the *P. vivax*/*An. stephensi* pair are similar to those for *P. falciparum*/*An. stephensi*. Again *µ* dominates at intermediate temperatures, but now near the upper thermal limit the uncertainty is almost entirely determined by uncertainty in vector competence (*bc*). At the lower limit, PDR contributed to the uncertainty but to a lesser extent than *µ* (See SM; Appendix B5d).

The patterns exhibited for suitability by *An. gambiae* are markedly different, reflecting the greater uncertainty across multiple traits used to construct the suitability metric. Although the uncertainty due to *µ* is the dominant source of uncertainty at intermediate temperatures, other parameters contribute to the uncertainty across much wider portions of the thermal response compared to *An. stephensi* (See SM; Appendix B6). For example, uncertainty for *P. falciparum*/*An. gambiae* pair is dominated by parasite development rate (PDR) in the lower and upper limits while eggs per female per day (EFD) and mosquito development rate (MDR) influenced uncertainty in the mid-to lower temperature ranges (See SM; Appendix B6c). For the *P. vivax*/*An. gambiae* pair, uncertainty near the lower and upper limits for transmission is dominated by EFD, MDR, and PDR, with each leading over slightly different ranges (See SM; Appendix B6d).

### 3.4 Model Validation

We assessed whether our suitability metric is consistent with patterns spatially explicit prevalence data for *P. falciparum* in Africa and Asia and *P. vivax* in Asia in two ways. For each set of parasite prevalence data we evaluated seven models that included a combination of the logarithm of population density, logarithm of GDP, and the probability that the suitability metric is greater than zero (i.e., *S*(*T*) > 0, SGTZ). This included three “null” models that incorporated socio-economic factors but not the suitability metric. We chose as the best model the one that has the lowest value of the Bayesian Information Criterion (BIC), and also evaluated the relative model probability based on BIC. We also used a visual, qualitative approach, that examines histograms of the proportion of *P. falciparum* and *P. vivax* positive prevalence cases that falls within suitable areas for malaria transmission (0 to 12 months) by *An. gambiae* and *An. stephensi* mosquitoes in Africa and Asia. (see SM; Appendices B19 and B20).

For *P. falciparum* in Africa and Asia the best model is the one that includes the linear combination of SGTZ, log population, and log GDP interacted with location (i.e., different regression coefficients for GDP in Africa than in Asia) (See SM; Appendix A9). This indicates that including the suitability metric is significantly better at explaining patterns of presence/absence than socio-economic factors alone. Based on the histograms of proportion of positive cases, we find that in Africa, 76% and 52% of *P. falciparum* positive prevalence have permissive temperatures ranges for at least six months of the year in areas suitable for *An. stephensi* and *An. gambiae* respectively. In Asia, 85% and 70% of *P. falciparum* positive prevalence have permissive temperatures ranges for at least six months of the year in areas suitable for *An. stephensi* and *An. gambiae* respectively. Together, these results indicate that our suitability metric is consistent with the observed patterns for *P. falciparum*.

In contrast, for *P. vivax* in Asia, when comparing the models based in BIC, the best model is the one that only includes the socio-economic factors. The second best model is the one that includes the linear combination of SGTZ with the socio-economic factors. The difference in BIC between these two best models is less than 10 units meaning that the difference in their performance are not highly significant (See SM; Appendix A10). The proportion of deviance explained by these models is 0.025, which is the same for both models. We expect that there are two primary reasons why we observe this. First, the overall amount of data is much smaller, which makes it more difficult to discern signals. Further, in the parts of Asia where we have data, temperature does not vary very much, where as there are larger swings in per capita GDP and population density; this is the opposite to the pattern observed in Africa (See SM; Appendix B21). Based on the histograms of proportion of positive cases, we find that in Asia, 69% and 70% of *P. vivax* positive prevalence cases have permissive temperature ranges for at least six months of the year in areas suitable for *An. stephensi* and *An. gambiae* respectively. In Africa, 38% and 52% of *P. vivax* positive prevalence have permissive temperatures ranges for at least six months of the year in areas suitable for *An. stephensi* and *An. gambiae* respectively (See SM; Appendices B19 and B20). This indicates that the suitability metrics are also likely consistent with the observed pattern of prevalence in both places, although further data to confirm this would be helpful.

### 3.5 Mapping climate suitability for malaria transmission

Our maps illustrate the number of months of suitability for *P. falciparum* and *P. vivax* transmission by *An. stephensi* and *An. gambiae* in Africa and for *P. falciparum* and *P. vivax* transmission by *An. stephensi* in Asia (Figure 4 and Appendix B18 in SM) based on the top quartile of the posterior probability median *S*(*T*) > 0. The maps predict the seasonality of temperature suitable for malaria transmission geographically, but they do not indicate its magnitude. Our maps demonstrate that temperatures are suitable for the transmission of *P. falciparum* and *P. vivax* malaria by both mosquito species, *An. gambiae* and *An. stephensi*, in vast areas of Africa and by *An. stephensi* mosquitoes in Asia (Figure 4 and Appendix B18 in SM) Our maps indicate that in Africa, approximately 15% of the continental land area will have temperatures suitable year-round for the transmission of *P. falciparum* malaria by *An. Stephensi* mosquitoes, and 44% for at least six months of the year (Figure 4A). We found that approximately 12% of the African continent is suitable for transmission of *P. vivax* by *An. stephensi* mosquitoes year-round, and 38% for at least six months of the year (Figure 4B). Countries in which most of the territory is suitable year-round include Liberia, Sierra Leone, Central African Republic, Democratic Republic of the Congo, Congo, Gabon, Cameroon, Cote d’Ivoire, and Ghana. The area suitable for *P. falciparum* and *P. vivax* malaria transmission by *An. gambiae* in Africa is very similar with 8% of the area suitable year-round and 30% of the area suitable at least six months of the year for both parasites (See SM; Appendix B18). Countries with most of their territory suitable year-round include Liberia, Democratic Republic of Congo, Congo, Central African Republic, and Cote d’Ivoire.

**Figure 4:**
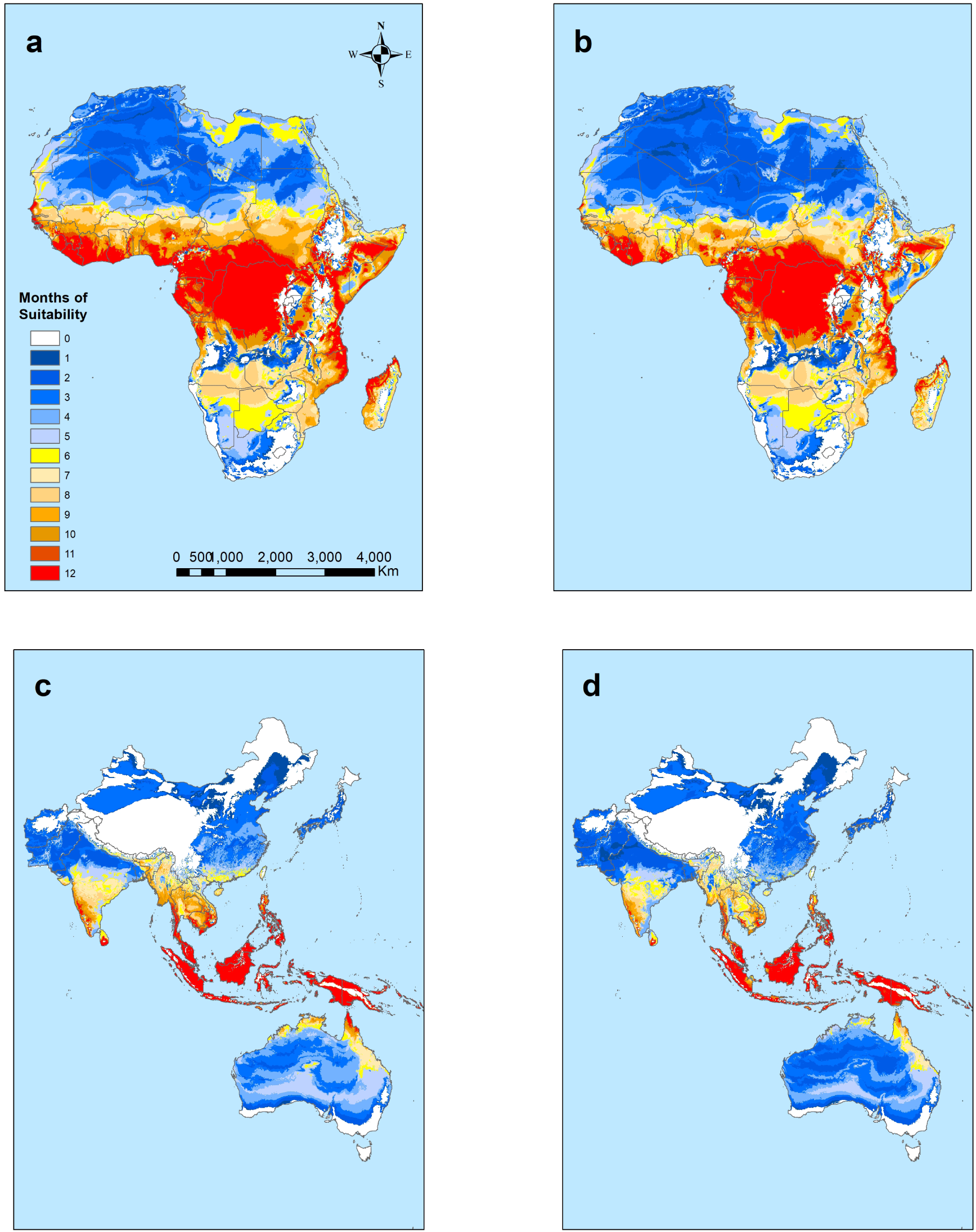
The number of months a year that locations in Africa and Asia are suitable (*S*(*T*)>0) for the transmission of A) *P. falciparum*, B) *P. vivax*, C) *P. falciparum* and, D) *P. vivax* by *An. stephensi* mosquitoes. We define highly suitable temperatures as *S*(*T*)>0.75.

In Asia, approximately 9% of the area is suitable for the transmission of *P. falciparum* malaria by *An. stephensi* mosquitoes year-round, and 20% for six or more months of the year (Figure 4C). The area suitable for year-round transmission of *P. vivax* by *An. stephensi* is approximately 7%, and for at least six months of the year is 16% of the Asian territory ((Figure 4D). Countries with most of their territory suitable year-round include Indonesia, Malaysia, Singapore, Papua New Guinea, the South of Thailand, and the South of Sri Lanka.

## 4 Discussion

Determining the optimal and the minimum and maximum temperature limits at which *An. gambiae* and *An. stephensi* mosquitoes are the most efficient vectors for the transmission of malaria parasites is important for assessing the potential for invasion and establishment in novel locations, and to predict the impacts of climate change on the future geographical distribution of these two mosquito species and the malaria parasites they transmit. In this paper, we have updated predictions for thermal suitability of transmission of the two most common malaria parasites, *P. falciparum* and *P. vivax*, by two of the most common malaria vector species, *An*.*gambiae* and *An. stephensi*, using data from historical and newly published studies. In doing so, we are able to examine the extent to which predictions may vary between mosquito species and between parasite types. We also identified persistent data gaps that must be addressed to further improve these models and allow more precise comparisons between mosquito/parasite complexes.

Our results suggest that there is little difference in the optimal temperature for malaria transmission between *An. gambiae* and *An. stephensi* mosquitoes. Further we find optimal temperatures for malaria transmission suitability that are similar to other recent findings for the optimal temperature at the continental scales (e.g., Johnson et al., 2015; Lunde et al., 2013; Mordecai et al., 2019, 2013; Shapiro et al., 2017). Earlier studies had calculated higher optimal temperatures for malaria transmission (Craig et al., 1999; Mahmood, 1997; Parham and Michael, 2009). We attribute the difference to the use of linearly increasing/monotonic functions as a component of the models. The study of (Shapiro et al., 2017) also reported an optimum temperature of 29 °C using a relative vectorial capacity model. Empirical data usually violates several of the vectorial capacity model assumptions (Shapiro et al., 2017). In nature, biological and ecological mosquito and parasite traits usually show unimodal responses to temperature (Dell et al., 2011). Biological and ecological traits in response to temperature tend to increase exponentially from a minimum thermal limit to an optimal temperature, then decline to a zero at a maximum thermal limit (Dell et al., 2011).

In contrast, we find that there are differences between the temperatures that could limit suitability for the two focal parasites to be spread by different mosquitoes. For example we found evidence that there are differences in the upper thermal limit between *P. vivax* and *P. falciparum* when spread by *An. stephensi*, and between *P. falciparum* when spread by the two mosquitoes. There is also evidence that the lower suitability threshold differs by mosquito species when transmitting the same pathogen. However, there is a great deal of uncertainty in the estimates of these limits due to poor data availability, especially for traits of *An. gambiae* and for *P. falciparum*.

As a result of the observed difference in the upper and lower thermal limits across mosquito-parasite systems, we also observe differences in the predicted temperature ranges for suitability, with greater temperature ranges for pathogens transmitted by *An. stephensi* than *An. gambiae* (Figure 3). Based on available trait data and previous studies, *An. stephensi* mosquitoes seem to have greater plasticity in thermal tolerance than *An. gambiae* mosquitoes (Bayoh, 2001; Kirby and Lindsay, 2004; Miazgowicz et al., 2020). Our predictions largely agree with these broader mosquito thermal studies.

The larger thermal breadth for transmission of both *P. falciparum* and *P. vivax* by *An. stephensi* than *An. gambiae* has a knock-on effect for the spatial extent of suitability for transmission. Our maps show the areas that are potentially currently suitable, based on temperature, for the transmission of *P. falciparum* and *P. vivax* by *An. gambiae* and *An. stephensi* mosquitoes. In Africa, vast regions between 22 °N and 21 °S are highly suitable for *An. gambiae* and *An. stephensi* mosquitoes. *An. gambiae* is currently the dominant mosquito species that transmits malaria in Africa. Although *An. stephensi* is a native mosquito of Asia (Sinka et al., 2010), recent research has reported that *An. stephensi* mosquitoes are already present in Africa, for example in Djibouti (Faulde et al., 2014), Ethiopia (Balkew et al., 2019; Carter et al., 2018), Sudan, and probably in neighboring countries (Takken and Lindsay, 2019; World Health Organization, 2019). The current presence and the possible spread of *An. stephensi* to African countries poses a potential health risk since it is a malaria vector well adapted to urban centers and it could cause malaria outbreaks of unprecedented sizes (Balkew et al., 2019; Takken and Lindsay, 2019). Our model indicates that the breadth of temperature range for *An. stephensi* with *P. falciparum* (15.3 to 37.2 °C) is greater than the breadth for *An. gambiae* with *P. falciparum* (19.1 to 30.1 °C); and in a lesser degree, the breadth for *An. stephensi* with *P. vivax* (15.7 to 32.5 °C) is greater than the breadth for *An. gambiae* with *P. vivax* (19.2 to 31.7 °C) (See SM; Appendix A7). This indicates that a larger proportion of Africa may be suitable for transmission by *An. stephensi* than by *An. gambiae*, due to the larger thermal breadth. Thus, regions at the northern and southern limits of the area dominated by *An. gambiae* are suitable for *An. stephensi*. In these areas malaria transmission could increase as *An. stephensi* becomes more established, and become a potential threat to malaria control in Africa.

Previous research assessing the optimal temperature and temperature limits for malaria transmission using mechanistic trait-based models has necessarily relied on data from a combination of mosquito and parasite species due to incomplete data availability for thermal traits of mosquitoes and parasites. In this study, all mosquito traits used for the calculations were from the *Anopheles* genus. If data for a specific trait were not available for one of the studied species, we used data from the closest relative in the same genus (see SM; Appendix A2), For example, for *An. stephensi*, all mosquito traits were from the same species, but for *An. gambiae* bite rate data are still not available so we used data from *An. arabiensis* and *An. pseudopunctipennis* mosquitoes instead. A similar approach was used for parasite traits. The amount of available data for each species has significant impact on the uncertainty in our model. To reduce uncertainty in these models, there is a need for more empirical data from the laboratory on how changes in temperature affect mosquito and parasite traits, especially for *An. gambiae*.For example, fecundity data for *An. gambiae* is lacking at low and high temperature ranges. There is also a need for vector competence and parasite development rate data for *An. stephensi* with *P. vivax, An. gambiae* with *P. vivax*, and *An. gambiae* with *P. falciparum*; and vector competence for *An. stephensi* with *P. falciparum*. These data would improve certainty in these models, especially at the thermal limits.

Our approach has some important limitations, some of which could be addressed by extending the mechanistic models. One limitation is that we use constant temperatures in our models, and did not incorporate daily and seasonal temperature variations, which occur in nature. However, non-linearities make it difficult to measure mosquito and parasite traits even at constant temperatures, especially at the thermal limits (Johnson et al., 2015; Mordecai et al., 2019). Precipitation influences malaria transmission, particularly for abundance, due to its role in mosquito life cycles. Incorporating the effect of different precipitation regimes on mosquito and parasites traits could also improve our mechanistic models. Despite these limitations, validation of these constant temperature mechanistic models, using prevalence data (our study), human case data (Mordecai et al., 2017), or entomological inoculation rate (EIR) data (Mordecai et al., 2013) demonstrate that these simple temperature-only models capture broad-scale patterns of transmission of mosquito-borne diseases. Due to the relative simplicity of the approach, similar studies combining empirical data and model fitting could estimate optimum temperature and the thermal limits for other vector-pathogen transmission systems in a similar way.

## 5 Acknowledgments

The authors want to thank Kerri Miazgowicz, Krijn Paaijmans and Antoine Barreaux for kindly providing raw data used in our analyses. LRJ, OVC, and SJR were supported by NSF EEID #1518681. LRJ was also supported by NSF DMS/DEB #1750113. CCM was supported by NIAID RO1 (1R01AI110793-01A1)

## Supplementary Appendices for

### A Supp Mat: Tables

#### A.1 Data sources and Species Information

**Table A1:** Available data on life history traits of mosquitoes that transmit malaria.

| Species | a | MDR | PEA | EFD | mu | bc | PDR | References |
| --- | --- | --- | --- | --- | --- | --- | --- | --- |
| *An. gambiae* |  | X | X | X | X |  | *P. falciparum* | (Afrane et al., 2008; Bayoh and Lindsay, 2003; Bayoh, 2001; Christiansen-Jucht et al., 2015; Kirby and Lindsay, 2009; Murdock et al., 2016; Olayemi and Ande, 2009) |
|  |  |  |  |  |  | X | *P. berghei* | (Barreaux et al., 2016) |
| *An. funestus* |  | X | X |  |  |  |  | (Lyons et al., 2013) |
| *An. arabiensis* | X | X | X | X |  |  |  | (Afrane et al., 2007; Kirby and Lindsay, 2009; Lyons et al., 2013; Maharaj, 1996) |
| *An. stephensi* | X | X | X | X | X | X | *P. falciparum* | (Eling et al., 2001; Miazgowicz et al., 2020; Murdock et al., 2016; Paaijmans et al., 2013a,b; Shapiro et al., 2017; Thomas et al., 2018) |
|  |  |  |  |  |  |  | *P. vivax* | (Thomas et al., 2018) |
|  |  |  |  |  |  |  | *P. malariae* | (Knowles et al., 1943) |
|  |  |  |  |  |  |  | *P. yoelii* | (Paaijmans et al., 2011, 2013a) |
| *An. quadrimaculatus* |  |  |  |  |  | X | *P. vivax* | (Boyd et al., 1932; Love and Whelchel, 1957; Stratman-Thomas, 1940) |
|  |  |  |  |  |  |  | *P. malariae* | (Boyd and Stratman-Thomas, 1933) |
| *An. culicifacies* |  |  |  |  |  |  | *P. malariae* | (Siddons et al., 1944) |
| *An. maculipennies* |  |  |  |  |  |  | *P. malariae* | (Shute and Maryon, 1952) |
| *An. pseudopunctipennies* | X |  |  |  |  |  |  | (Lardeux et al., 2008) |
| *An. albimanus* | X | X | X |  |  |  | *P. falciparum* | (Grieco et al., 2003; Mahmood, 1997; Vélez, 2008) |

**Table A2:** Ecological and biological comparison between mosquito species that transmit malaria

| Species | Feeding habit | Biting habit | Resting | Breeding | Genome Similarities | Geographic distribution | References |
| --- | --- | --- | --- | --- | --- | --- | --- |
| *An. gambiae* | Anthropophilic | Endophagic | Endophilic (exophilic*) | Shaded pools, rice fields, tolerates salty water | *An. arabiensis* | Africa | (Neafsey et al., 2015; O'Loughlin et al., 2014; Paaijmans and Thomas, 2011; Sinka et al., 2012) |
| *An. funestus* | Anthropophilic | Endophagic | Endophilic | Rice puddles, large ponds, lake edges | *An. stephensi*,<br>*An. culicifacies*,<br>*An. minimus* | Africa | (Carter et al., 2018; Paaijmans and Thomas, 2011; Sinka et al., 2012) |
| *An. arabiensis* | Zoophilic | Exophagic | Exophilic | Diverse natural and manmade places | *An. gambiae* | Africa | (Carter et al., 2018; Mahande et al., 2007; Paaijmans and Thomas, 2011; Sinka et al., 2012) |
| *An. stephensi* | Zoophilic | Exophagic | Exophilic | Manmade containers, overhead and, underground tanks | *An. funestus*,<br>*An. culicifacies*,<br>*An. maculipennis* | Western Asia, India, China, Ethiopia | (Carter et al., 2018; Paaijmans and Thomas, 2011; Sinka et al., 2012) |
| *An. quadrimaculatus* | Zoophilic and anthropophilic | Exophagic | Exophilic | Farm fields, lakes, swamps, old tires, animal stables | *An. maculipennis* | Mexico, Southeastern USA | (Neafsey et al., 2015; Rios and Connelly, 2012; Sinka et al., 2012) |
| *An. culicifacies* | Anthropophilic and zoophilic | Endophagic and exophagic | Endophilic | Irrigated channels, rock and sandy pools, borrow pits, rice fields | *An. stephensi*<br>*An. funestus* | India, China<br>Ethiopia | (Neafsey et al., 2015; Sharma and Dev, 2015; Sinka et al., 2012) |
| *An. maculipennis* | Zoophilic and anthropophilic | Exophagic | Exophilic (endophilic*) | Cannals, ditches, river margins, rice fields | *An. quadrimaculatus* | South-East Asia pacific, North-East Europe | (Neafsey et al., 2015; Ponçon et al., 2007; Sinka et al., 2012) |
| *An. pseudo-punctipennis* | Zoophilic and, anthropophilic | Exophagic and endophagic | Exophilic (endophilic*) | Lagoons, slow flow rivers |  | South and central America | (Lardeux et al., 2007; Sinka et al., 2011, 2010, 2012) |
| *An. albimanus* | Zoophilic | Exophagic | Exophilic (endophilic*) | Places with salty and brackish water |  | Neotropics (South and central America) | (Neafsey et al., 2015; Paaijmans and Thomas, 2011; Sinka et al., 2012) |

\* Presence of behaviour in rare occasions

#### A.2 Priors for Bayesian Analysis of Thermal Traits

**Table A3:** Prior distributions for each of the parameters for the best fitting of the responses for each of the thermal traits used for *S*(*T*) calculation.

| Thermal response function | Model parameters | Parameters | Priors |
| --- | --- | --- | --- |
| Brière | Mosquito development rate (MDR)<br>Parasite development rate (PDR)<br>Bite rate($a$) | $T_0$<br>$T_m$<br>$c$<br>$\tau$ | $dunif(0,24)$<br>$dunif(25,45)$<br>$dgamma(1,10)$<br>$dgamma(0.0001,0.0001)$ |
| Quadratic<br>(Concave down) | Eggs to adulthood survival ($P_{EA}$)<br>Eggs per female per day (EFD) | $T_0$<br>$T_m$<br>$qd$<br>$\tau$ | $dunif(0,24)$<br>$dunif(25,45)$<br>$dgamma(1,1)$<br>$dgamma(0.0001,0.0001)$ |
| Quadratic<br>(Concave down) | Vector competence (bc) | $T_0$<br>$T_m$<br>$n.qd$<br>$qd$ | $dunif(0,24)$<br>$dunif(25,45)$<br>$dexp(1)$<br>$-n.qd$ |
| Quadratic<br>(Concave up) | Mortality rate ($\mu$) | $int$<br>$n.slope$<br>$qd$<br>$\tau$ | $dgamma(2,2)$<br>$dgamma(1,1)$<br>$dgamma(2,2)$<br>$dnorm(1000,1/500)$ |

#### A.3 Mosquito and parasites parameters to calculate S(T)

**Table A4:**
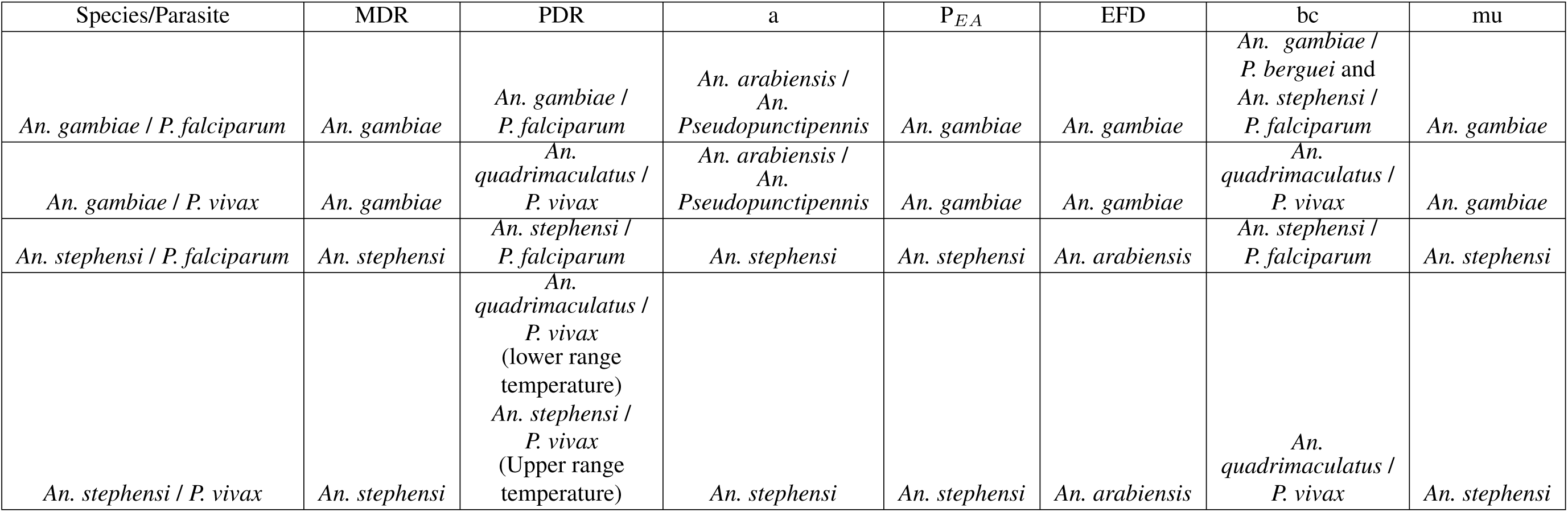
Mosquito and parasite parameters used to calculate the suitability matrix (*S*(*T*)) in *An. gambiae* and *An. stephensi* mosquitoes.

#### A.4 Summaries of Thermal Traits

**Table A5:** Mean and 95% credible intervals (95% CI) on the critical thermal minimum, maximum, and optimum temperature for mosquito and parasite traits for *An. gambiae* mosquitoes.

| *Anopheles gambiae* | Mean T peak<br>(95% CI) | Mean T min<br>(95% CI) | Mean T max<br>(95% CI) |
| --- | --- | --- | --- |
| Mosquito development rate | 29<br>(28.46 - 29.54) | 13.85<br>(13.02 - 14.68) | 35.95<br>(35.54 - 36.36) |
| Bite rate | 34.6<br>(33.89 - 35.31) | 16.95<br>(16.14 - 17.76) | 43.55<br>(43.24 - 43.86) |
| Eggs per female per day | 25.9<br>(25.37 - 26.43) | 15.30<br>(14.65 - 15.95) | 32.70<br>(32.27 - 33.13) |
| Proportion of eggs to adulthood | 24.80<br>(24.17 - 25.43) | 16.65<br>(16.27 - 17.03) | 32.70<br>(32.34 - 33.06) |
| Mortality rate | 22.30<br>(21.70 - 22.89) | 8.45<br>(7.70 - 9.19) | 38.85<br>(37.99 - 39.70) |
| Vector competence<br>*P. falciparum* | 24<br>(23.37 - 24.63) | 9.45<br>(8.70 - 10.19) | 36.15<br>(35.52 - 36.78) |
| Vector competence<br>*P. vivax* | 22.15<br>(21.61 - 22.69) | 13.55<br>(13.02 - 14.08) | 30.10<br>(29.61 - 30.59) |
| Parasite development rate<br>*P. falciparum* | 25.60<br>(25.06 - 26.14) | 12.35<br>(11.59 - 13.11) | 36.60<br>(35.93 - 37.26) |
| Parasite development rate<br>*P. vivax* | 28<br>(27.53 - 28.47) | 20<br>(19.44 - 20.56) | 31.65<br>(31.44 - 31.85) |

**Table A6:**
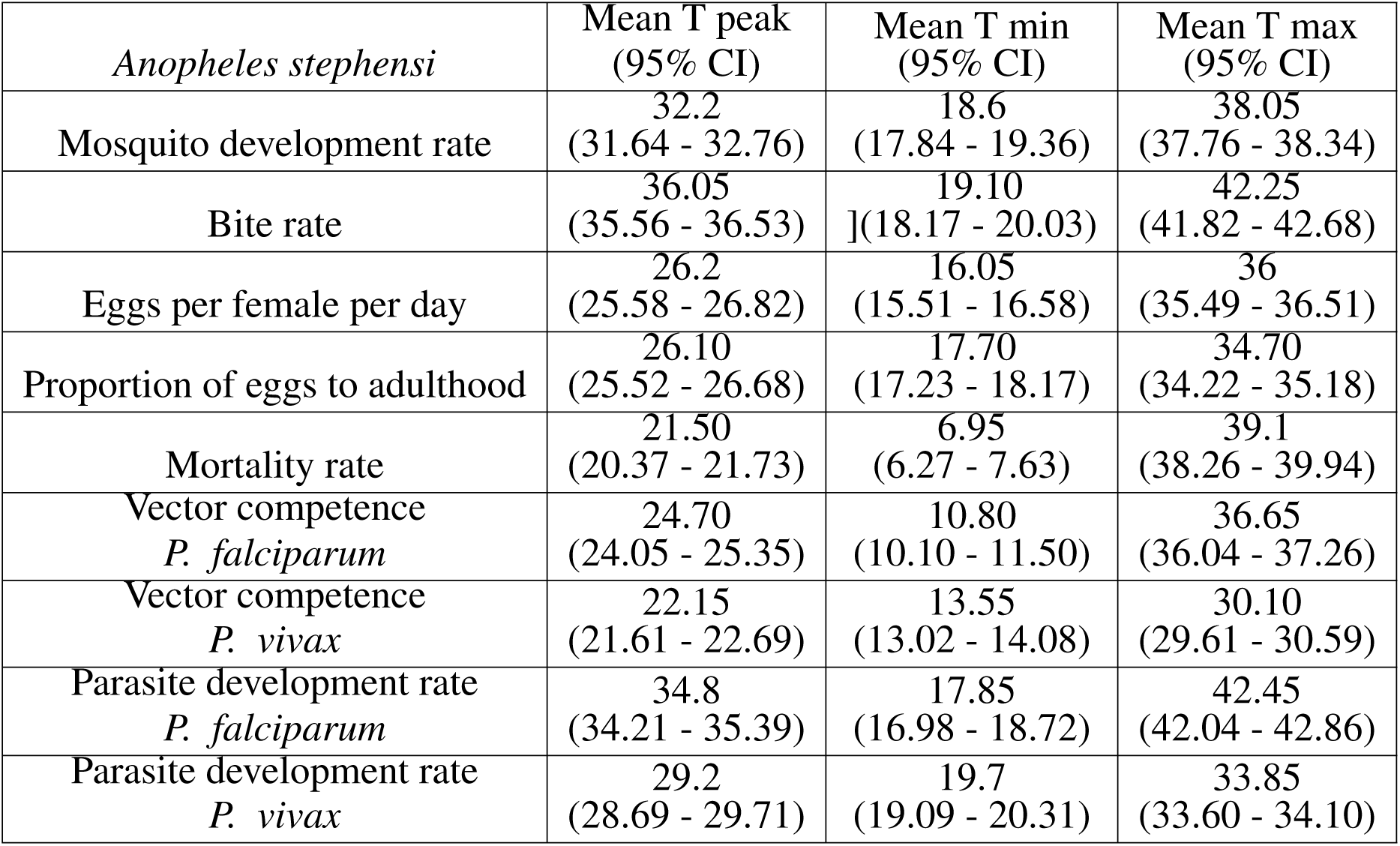
Mean and 95% credible intervals (95% CI) on the critical thermal minimum, maximum, and optimum temperature for mosquito and parasite traits for *An. stephensi* mosquitoes.

| *Anopheles stephensi* | Mean T peak<br>(95% CI) | Mean T min<br>(95% CI) | Mean T max<br>(95% CI) |
| --- | --- | --- | --- |
| Mosquito development rate | 32.2<br>(31.64 - 32.76) | 18.6<br>(17.84 - 19.36) | 38.05<br>(37.76 - 38.34) |
| Bite rate | 36.05<br>(35.56 - 36.53) | 19.10<br>(18.17 - 20.03) | 42.25<br>(41.82 - 42.68) |
| Eggs per female per day | 26.2<br>(25.58 - 26.82) | 16.05<br>(15.51 - 16.58) | 36<br>(35.49 - 36.51) |
| Proportion of eggs to adulthood | 26.10<br>(25.52 - 26.68) | 17.70<br>(17.23 - 18.17) | 34.70<br>(34.22 - 35.18) |
| Mortality rate | 21.50<br>(20.37 - 21.73) | 6.95<br>(6.27 - 7.63) | 39.1<br>(38.26 - 39.94) |
| Vector competence<br>*P. falciparum* | 24.70<br>(24.05 - 25.35) | 10.80<br>(10.10 - 11.50) | 36.65<br>(36.04 - 37.26) |
| Vector competence<br>*P. vivax* | 22.15<br>(21.61 - 22.69) | 13.55<br>(13.02 - 14.08) | 30.10<br>(29.61 - 30.59) |
| Parasite development rate<br>*P. falciparum* | 34.8<br>(34.21 - 35.39) | 17.85<br>(16.98 - 18.72) | 42.45<br>(42.04 - 42.86) |
| Parasite development rate<br>*P. vivax* | 29.2<br>(28.69 - 29.71) | 19.7<br>(19.09 - 20.31) | 33.85<br>(33.60 - 34.10) |

#### A.5 Summaries of the Suitability Metric, *S*(*T*)

**Table A7:** Mean and 95% credible intervals (95% CI) on the critical thermal minimum, maximum, and optimum temperature for suitability of malaria by *An. stephensi* and *An. gambiae* mosquitoes, based on the posterior distribution of *S*(*T*)

| Species/Parasite | Mean T peak<br>(95% CI) | Mean T min<br>(95% CI) | Mean T max<br>(95% CI) |
| --- | --- | --- | --- |
| *An. stephensi*/*P. falciparum* | 24.8<br>(23.2 - 26.1) | 15.3<br>(14.6 - 16) | 37.2<br>(36.5 - 38) |
| *An. stephensi*/*P. vivax* | 24.5<br>(23.2 - 25.6) | 15.7<br>(14.9 - 16.6) | 32.5<br>(31.7 - 33.7) |
| *An. gambiae*/*P. falciparum* | 25<br>(23.7 - 26.1) | 19.1<br>(15.5 - 22.5) | 30.1<br>(27.4 - 33.5) |
| *An. gambiae*/*P. vivax* | 25<br>(23.8 - 26) | 19.2<br>(16.6 - 22) | 31.7<br>(30.4 - 32.1) |

#### A.6 Posterior samples differences between mosquito/parasite systems

**Table A8:** Comparison of the samples of the *S*(*T*) posterior distributions at the optimum temperature, and maximum and minimum temperature limit. Values are expressed as proportions

| Sample differences | >0 | <0 |
| --- | --- | --- |
| Species/Parasite | Maximum temperature limit |  |
| *An. stephensi* - *P. falciparum* vs<br>*An. stephensi* - *P. vivax* | 1.00 | 0.00 |
| *An. gambiae* - *P. falciparum* vs<br>*An. gambiae* - *P. vivax* | 0.27 | 0.71 |
| *An. stephensi* - *P. falciparum* vs<br>*An. gambiae* - *P. falciparum* | 1.00 | 0.00 |
| *An. stephensi* - *P. vivax* vs<br>*An. gambiae* - *P. vivax* | 0.91 | 0.05 |
|  | Optimum temperature limit |  |
| *An. stephensi* - *P. falciparum* vs<br>*An. stephensi* - *P. vivax* | 0.61 | 0.36 |
| *An. gambiae* - *P. falciparum* vs<br>*An. gambiae* - *P. vivax* | 0.46 | 0.49 |
| *An. stephensi* - *P. falciparum* vs<br>*An. gambiae* - *P. falciparum* | 0.44 | 0.51 |
| *An. stephensi* - *P. vivax* vs<br>*An. gambiae* - *P. vivax* | 0.28 | 0.67 |
|  | Minimum temperature limit |  |
| *An. stephensi* - *P. falciparum* vs<br>*An. stephensi* - *P. vivax* | 0.21 | 0.73 |
| *An. gambiae* - *P. falciparum* vs<br>*An. gambiae* - *P. vivax* | 0.47 | 0.51 |
| *An. stephensi* - *P. falciparum* vs<br>*An. gambiae* - *P. falciparum* | 0.00 | 1.00 |
| *An. stephensi* - *P. vivax* vs<br>*An. gambiae* - *P. vivax* | 0.01 | 0.99 |

#### A.7 Model Validation Results

**Table A9:** Logistic regression models used for validation exercise – to test if the probability *S*(*T*)>0 alone or in combination with socio-economic variables is a good predictor of malaria prevalence caused by *P. falciparum* in Africa and Asia. For Africa and Asia we have 8343 and 3001 prevalence records respectively with 7934 *P. falciparum* positive prevalence cases and 3410 *P. falciparum* negative prevalence cases. The logistic model is defined so that for each response, *y*_*i*_, *Pr*(*y*_*i*_ = 1) = *θ* = logit^−1^(*η*_*i*_) where *η*_*i*_ is the linear predictor as given in the table above. Observations at data point *i* are then Binomial random variables with “success” probability *θ* and sample size *n*_*i*_. Abbreviations used on the table are explained below.

| Model | Formula for linear predictor | BIC | Model pr. | $D^2$ |
| --- | --- | --- | --- | --- |
| *Pfal_PA1* | $\eta_i \sim \beta_0 + \beta_1 lpop\_den + \beta_2 lpop\_den \mathbb{1}_{AFR}$ | 13175.29 | 0 | 0.052 |
| *Pfal_PA2* | $\eta_i \sim \beta_0 + \beta_1 lGDP + \beta_2 lGDP \mathbb{1}_{AFR}$ | 12984.26 | 0 | 0.066 |
| *Pfal_PA3* | $\eta_i \sim \beta_0 + \beta_1 SGTZ + \beta_2 SGTZ \mathbb{1}_{AFR}$ | 13072.04 | 0 | 0.060 |
| *Pfal_PA4* | $\eta_i \sim \beta_0 + \beta_1 lpop\_den + \beta_2 lGDP + (\beta_3 lpop\_den + \beta_4 lGDP) \mathbb{1}_{AFR}$ | 12933.72 | 0 | 0.071 |
| *Pfal_PA5* | $\eta_i \sim \beta_0 + \beta_1 SGTZ + \beta_2 lpop\_den + \beta_3 lGDP + (\beta_4 SGTZ + \beta_5 lpop\_den + \beta_6 lGDP) \mathbb{1}_{AFR}$ | 12658.45 | 1 | 0.092 |
| *Pfal_PA6* | $\eta_i \sim \beta_0 + \beta_1 \log(SGTZ \times pop\_den) + \beta_2 \log(SGTZ \times pop\_den) \mathbb{1}_{AFR}$ | 13023.10 | 0 | 0.063 |
| *Pfal_PA7* | $\eta_i \sim \beta_0 + \beta_1 \log(SGTZ \times GDP) + \beta_2 \log(SGTZ \times GDP) \mathbb{1}_{AFR}$ | 12776.25 | 0 | 0.081 |

**Table A10:** Logistic regression models used for validation exercise – to test if the probability of *S*(*T*)>0 alone or in combination with socio-economic is a good predictor of malaria prevalence caused by *P. vivax* in Asia. For Asia we have 3112 prevalence records with 1386 *P. vivax* positive prevalences cases and 1726 negative *P. vivax* prevalence cases. For Africa, there is not available *P. vivax* prevalence data. The logistic model is defined so that for each response, *y*_*i*_, *Pr*(*y*_*i*_ = 1) = *θ* = logit^−1^(*η*_*i*_) where *η*_*i*_ is the linear predictor as given in the table above. Observations at data point *i* are then Binomial random variables with “success” probability *θ* and sample size *n*_*i*_. Abbreviations used on the table are explained below.

| Model | Formula (linear predictor) | BIC | Model pr. | $D^2$ |
| --- | --- | --- | --- | --- |
| *Pviv_PA1* | $\eta_i \sim \beta_0 + \beta_1 lpop\_den$ | 4238.11 | 0 | 0.013 |
| *Pviv_PA2* | $\eta_i \sim \beta_0 + \beta_1 lGDP$ | 4223.61 | 0 | 0.016 |
| *Pviv_PA3* | $\eta_i \sim \beta_0 + \beta_1 SGTZ$ | 4284.74 | 0 | 0.002 |
| *Pviv_PA4* | $\eta_i \sim \beta_0 + \beta_1 lpop\_den + \beta_2 lGDP$ | 4193.86 | 0.98 | 0.025 |
| *Pviv_PA5* | $\eta_i \sim \beta_0 + \beta_1 SGTZ + \beta_2 lpop\_den + \beta_3 lGDP$ | 4201.72 | 0.02 | 0.025 |
| *Pviv_PA6* | $\eta_i \sim \beta_0 + \beta_1 \log(SGTZ \times pop\_den)$ | 4264.92 | 0 | 0.007 |
| *Pviv_PA7* | $\eta_i \sim \beta_0 + \beta_1 \log(SGTZ \times GDP)$ | 4266.28 | 0 | 0.006 |

### B Supp Mat: Figures

#### B.1 Posterior mean and 95% HPD of the thermal responses for *Anopheles gambiae* traits

**Figure B1:**
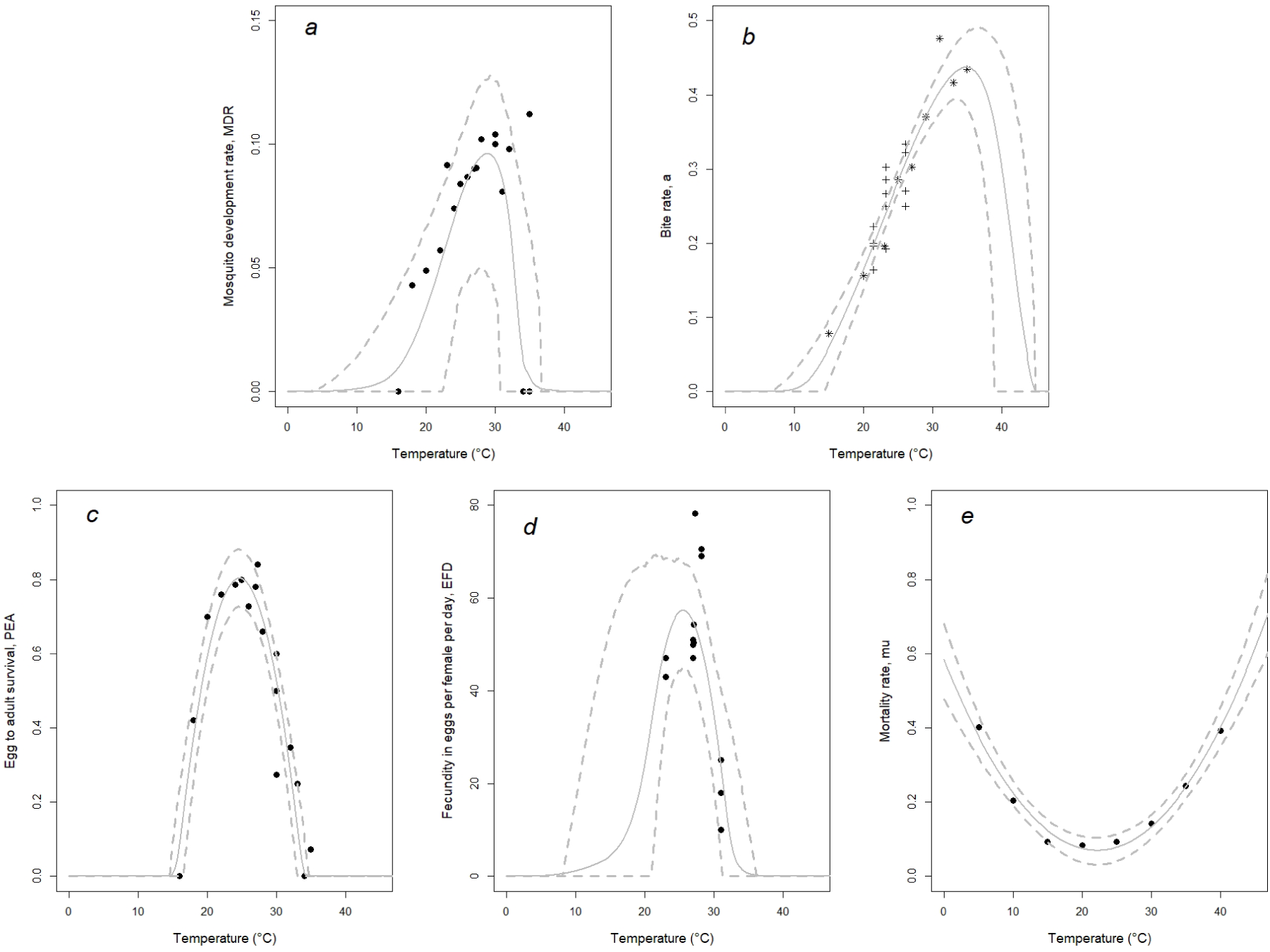
Posterior mean (solid line) and 95% highest posterior density-HPD (dashed lines) of the thermal responses for mosquito traits to calculate *S*(*T*) for *Anopheles gambiae*/*Plasmodium falciparum* and *An. gambiae*/*P. vivax* combinations. Traits modeled with a Briére thermal response are (A) mosquito development rate and (B) bite rate. Traits modeled with a concave down quadratic function are (C) proportion of eggs to adult and (D) fecundity; and (E) mortality rate is modeled with a concave-up quadratic function. Data symbols correspond to the species of mosquitoes and/or parasite used for the analysis. •: *An. gambaie*; + : *An. arabiensis*; ∗: *An. pseudopunctipennies*

### B.2 Posterior mean and 95% HPD of the thermal responses for *An. gambiae*/parasites traits

**Figure B2:**
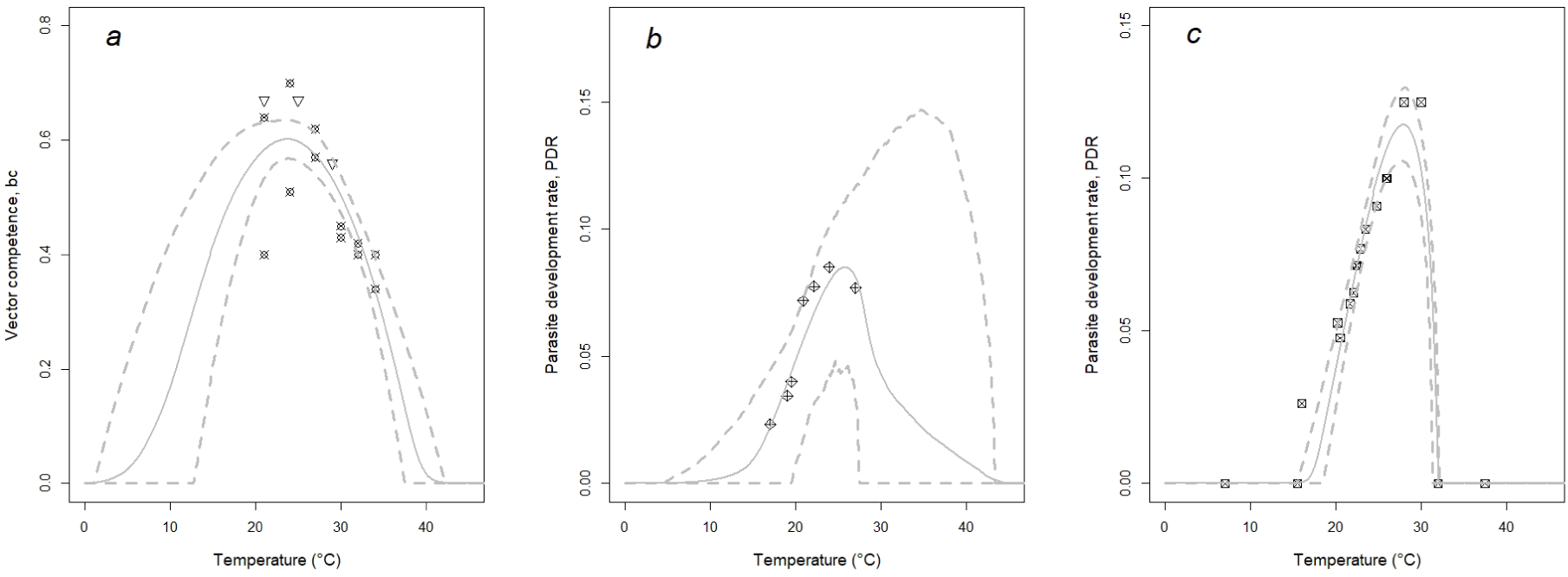
Posterior mean (solid line) and 95% highest posterior density-HPD (dashed lines) of the thermal responses for the mosquito/parasite traits to calculate *S*(*T*) for *Anopheles gambiae*/*Plasmodium faclciparum* and *An. gambiae*/*P. vivax*. Traits modeled with a concave down quadratic function are vector competence (A) *P. falciparum*. Traits modeled with a Briére thermal response are parasite development rates (B) *P. falciparum* (C) *P. vivax*. For Vector competence for *An. gambiae*/*P. vivax*, we used the same data as for *An. stephensi*/*P*.*vivax*. Data symbols correspond to the species of mosquito and parasite used for the analysis. ⊗ is for *An. stephensi* in combination with *P. falciparum*, ▽ is for *An. gambiae* in combination with *P. berghei*, ⊞ is for *An. gambiae* in combination with *P. falciparum*, and ⊠ is for *An. quadrimaculatus* in combination with *P. vivax*.

### B.3 Comparison of the thermal responses for parasite and mosquito traits

**Figure B4:**
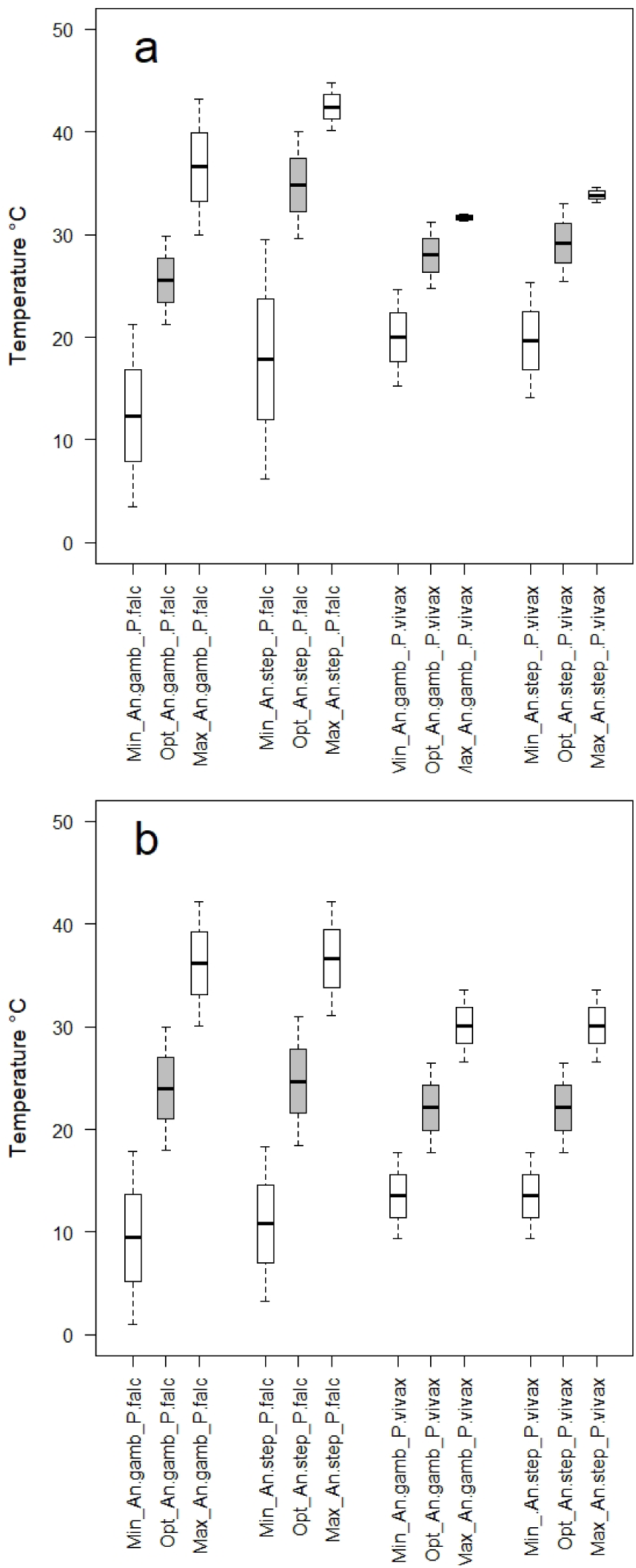
Comparison of the thermal response for parasites traits between mosquito/parasite systems: *An. gambiae*/*P. falciparum* (An. Gamb_P. falc), *An. gambiae*/*P. vivax* (An. Gamb_P. vivax), *An. stephensi*/*P. falciparum* (An. step P. falc), and *An. stephensi*/*P. vivax* (An. step P. vivax). A) Parasite development rate and B) Vector competence. Min: Minimum temperature, Opt: optimum temperature, and Max: Maximum temperature.

**Figure B3:**
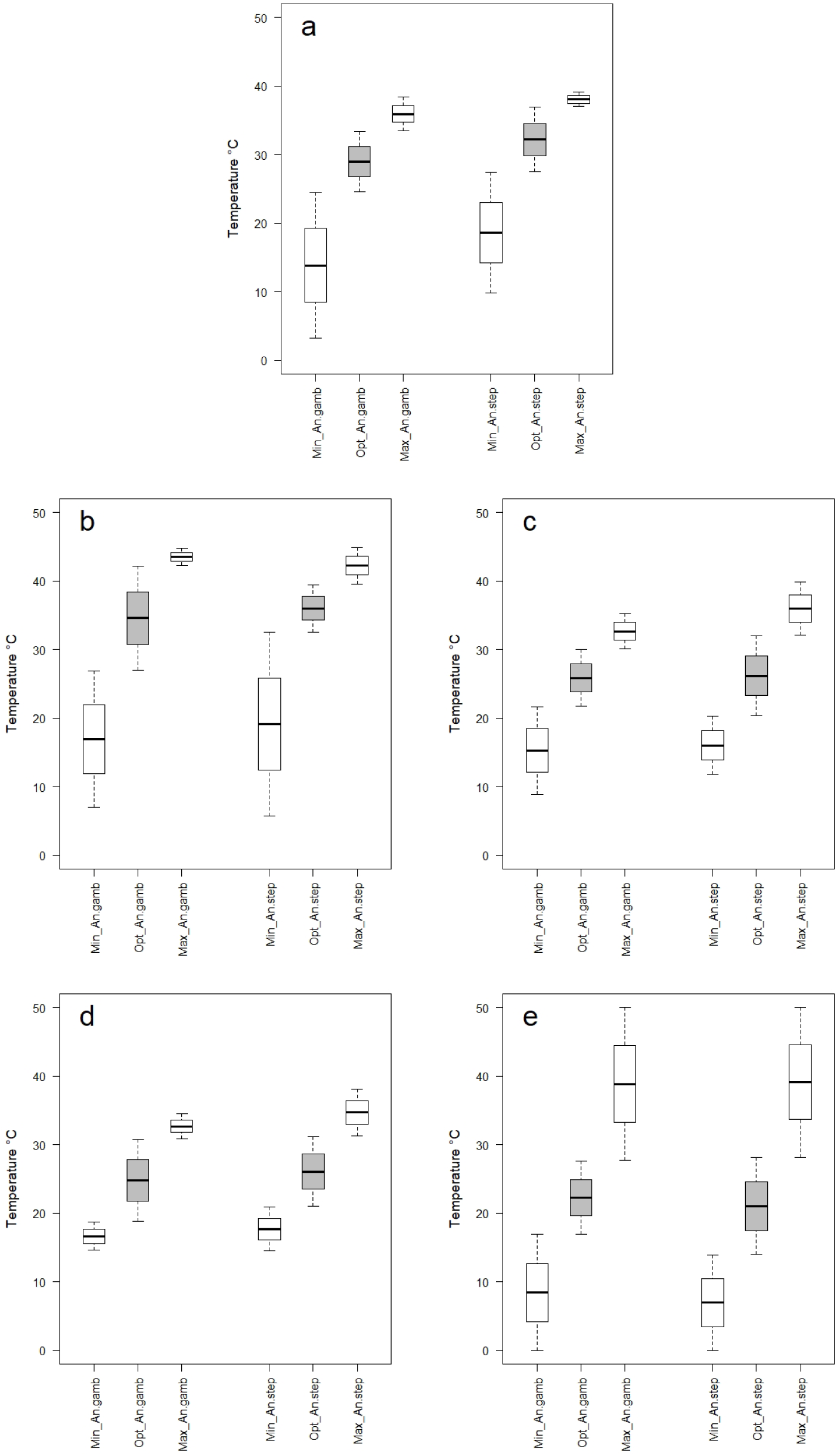
Comparison of the thermal responses for mosquito traits between *Anopheles gambiae* and *An. stephensi* mosquitos. (A) mosquito development rate, (B) bite rate (C) fecundity (D) proportion of eggs to adult and (E) mortality rate.

### B.4 Thermal response S(T) and its uncertainty for mosquito/parasite systems

**Figure B5:**
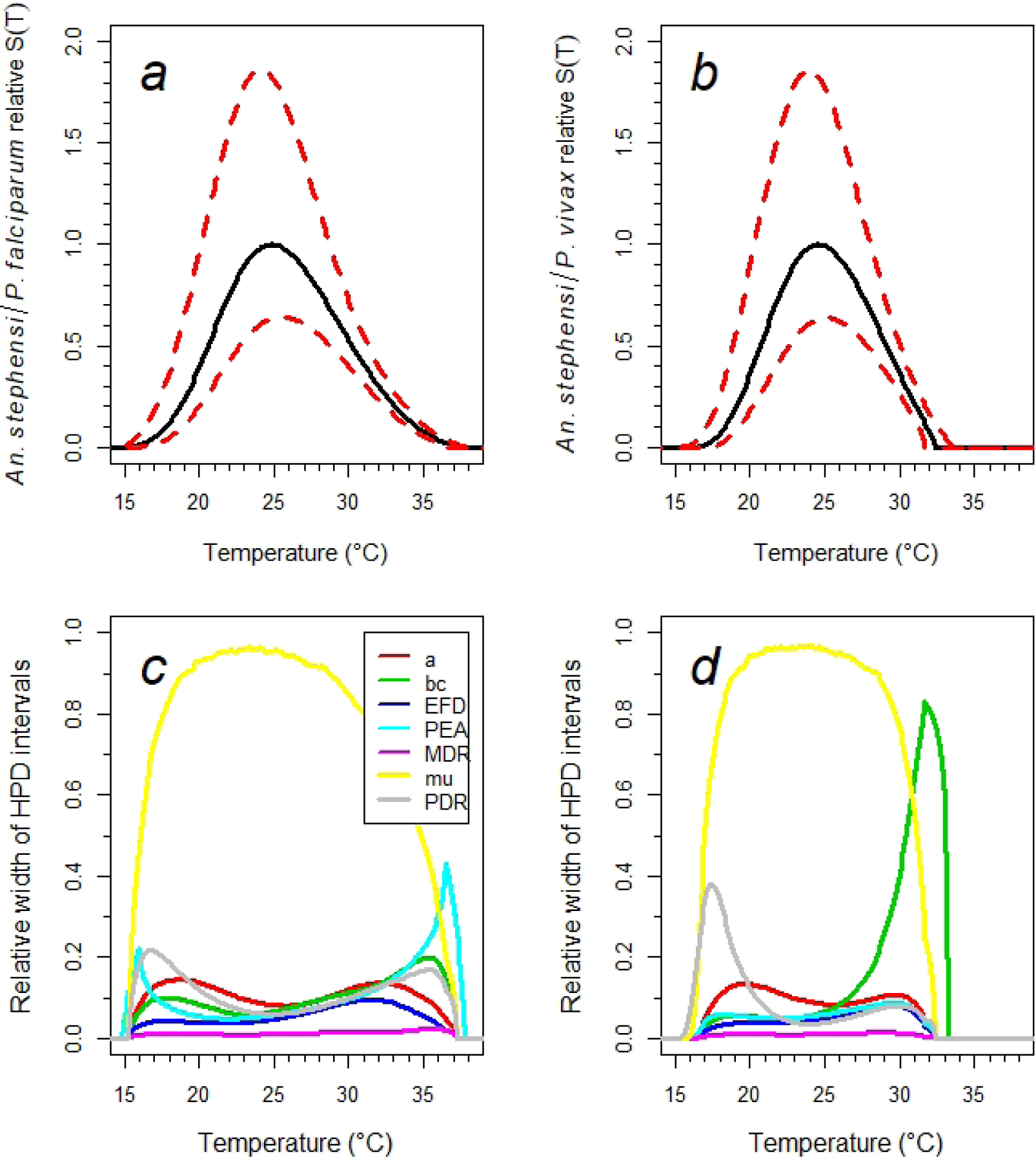
Relative *S*(*T*) divided by the maximum value of the posterior mean for A) *An. stephensi* with *P. falciparum* and B) *An. stephensi* with *P. vivax*. Relative width of the 95% HPD intervals due to uncertainty in each component, compared to uncertainty in *S*(*T*) overall for C) *An. stephensi* and *P. falciparum* and D) *An. stephensi* and *P. vivax*. Each curve was obtained as followed. For each component, *S*(*T*) was calculated for the thinned posterior samples of that component, with all other components set to its posterior mean. Then the width of the inner 95% HPD was calculated at each temperature. This was them normalized to the width of the HPD of the full posterior distribution of *S*(*T*) at each temperature

**Figure B6:**
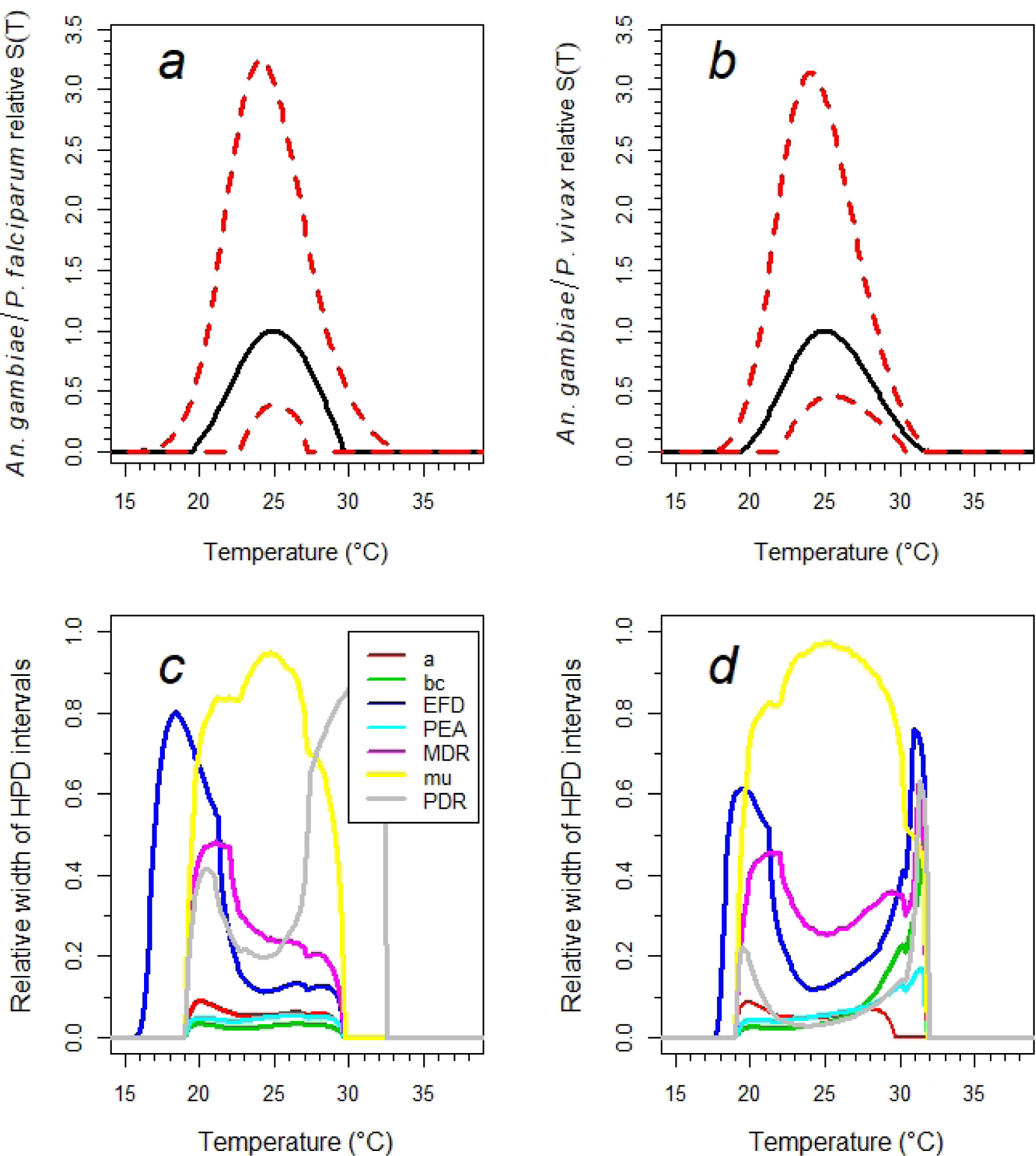
Relative *S*(*T*) divided by the maximum value of the posterior mean for A) *An. gambiae* with *P. falciparum* and B) *An. gambiae* with *P. vivax*. Relative width of the 95% HPD intervals due to uncertainty in each component, compared to uncertainty in *S*(*T*) overall for C) *An. gambiae* and *P. falciparum* and D) *An. gambiae* and *P. vivax*. Each curve was obtained as followed. For each component, *S*(*T*) was calculated for the thinned posterior samples of that component, with all other components set to its posterior mean. Then the width of the inner 95% HPD was calculated at each temperature. This was them normalized to the width of the HPD of the full posterior distribution of *S*(*T*) at each temperature

### B.5 Model validation

**Figure B7:**
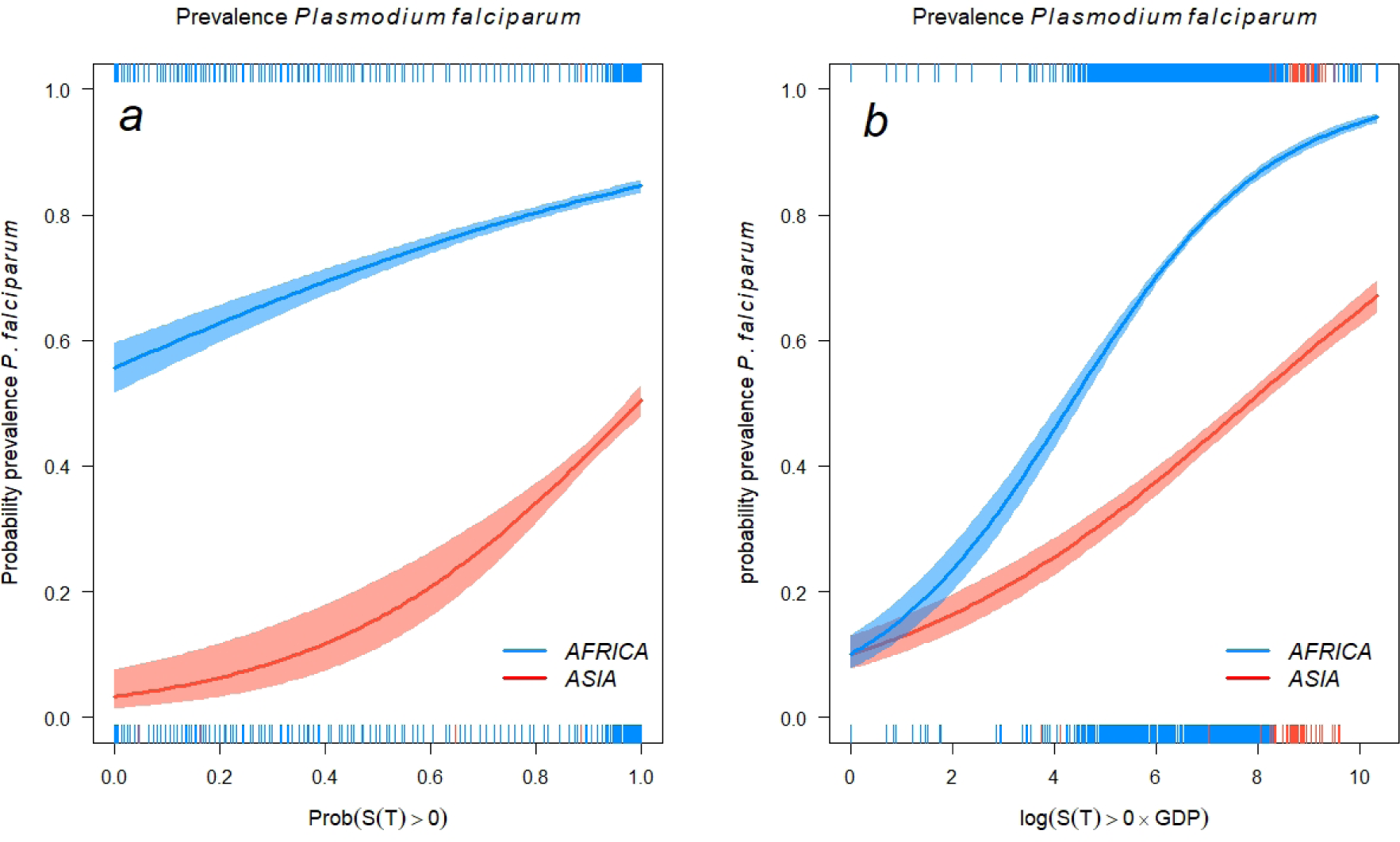
The probability of *S*(*T*)>0 and the per capita Gross Domestic Product (GDP) predict the probability of prevalence of *Plasmodium falciparum* in Africa and Asia. A) The posterior probability that *S*(*T*)>0 versus the probability of malaria prevalence caused by *P. falciparum* and transmitted by *An. gambiae* in Africa and by *An. stephensi* in Asia. B) the natural log of the probability of *S*(*T*) > 0 times the per capita gross domestic product versus the probability of malaria prevalence caused by *P. falciparum* and transmitted by *An. gambiae* in Africa and *An. stephensi* in Asia. For these models the median value of population density is 66.74 people/km^2^ and the median value of GDP is $1815.3. Tick-marks are positive and negative residuals on the top and bottom axes. Lines and shaded areas: mean and 95% CI from GLM fits for Africa (blue) and Asia (red). Figures for the probability prevalence caused by *P. vivax* and the interaction between *S*(*T*) > 0 and the socioeconomic variables are included in the supplemental material.

**Figure B8:**
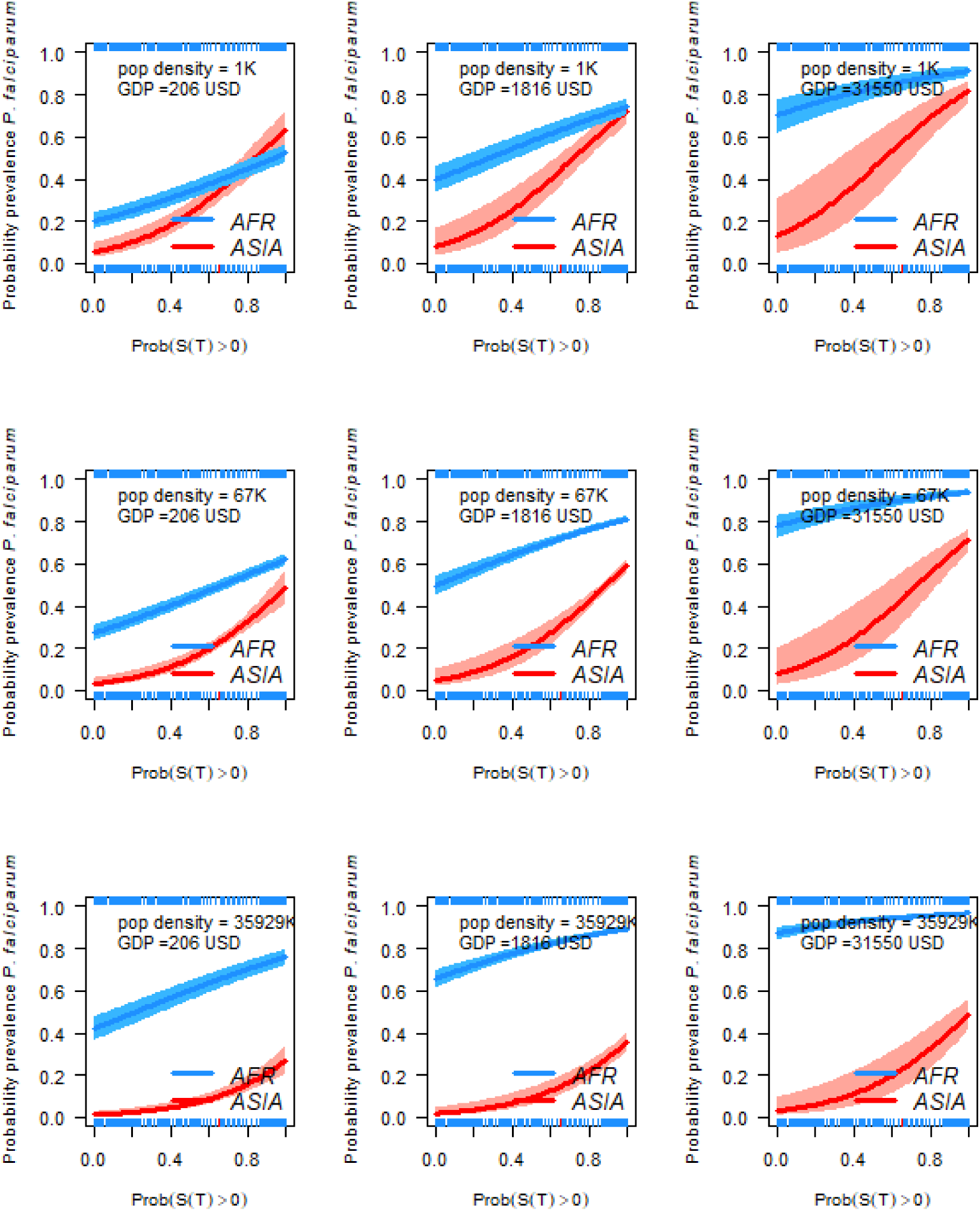
Plots of the probability that *S*(*T*) >0 versus the probability of malaria prevalence by *Plasmodium falciparum* transmitted by *An. gambiae* in Africa and by *An. stephensi* in Asia from presence/absence model *Pfalc* PA5 (Table 8; SM), for different levels of population density across different rows and different levels of per capita Gross Domestic Product (GDP) across different columns. Tick-marks are human malaria prevalence data. Lines and shaded areas: mean and 95% confidence interval from the fitted GLM.

**Figure B9:**
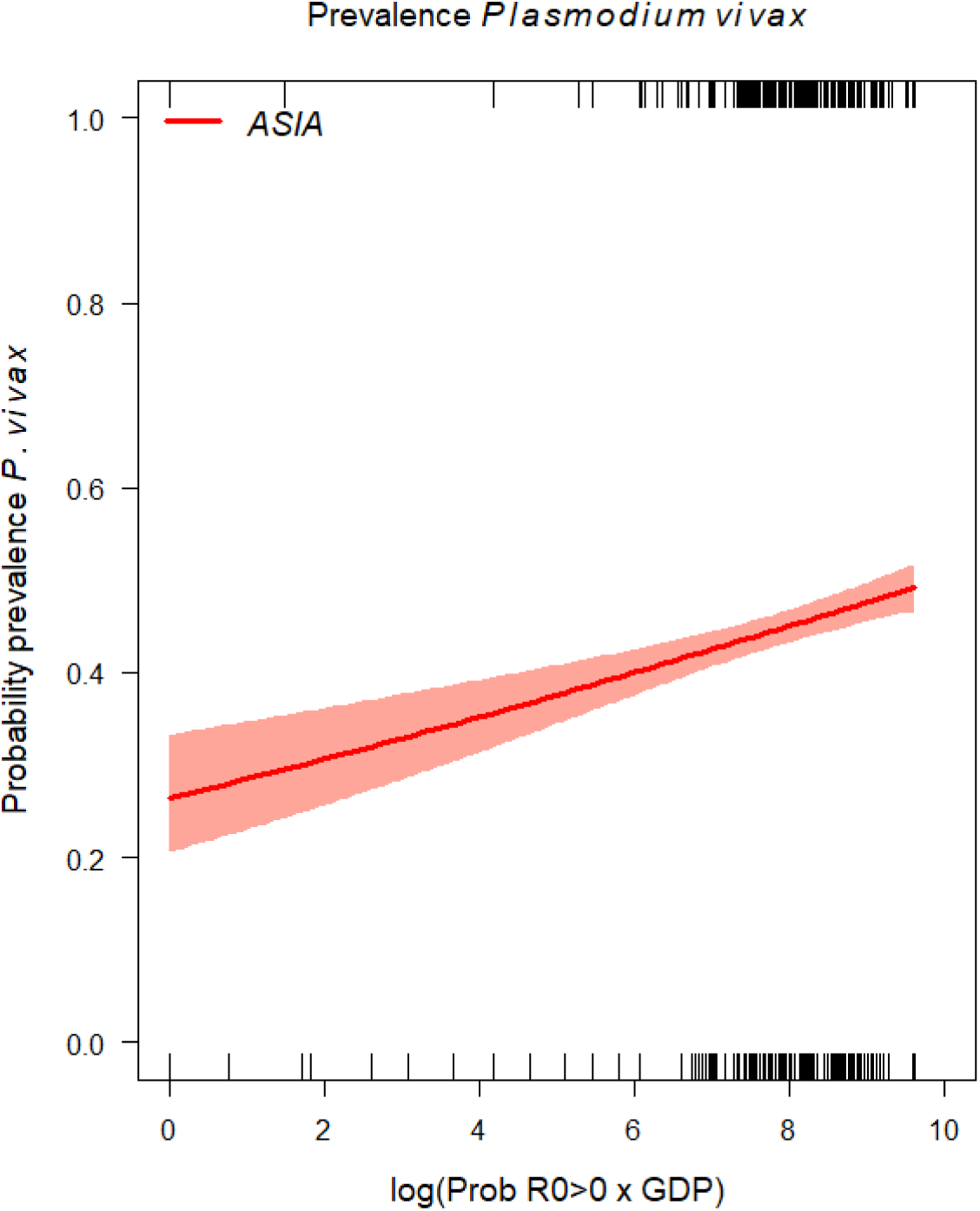
The natural log of the probability of *S*(*T*)>0 times the per capita Gross Domestic Product (GDP) predict the probability of prevalence of *P. vivax* transmitted by *An. stephensi* in Asia.

**Figure B10:**
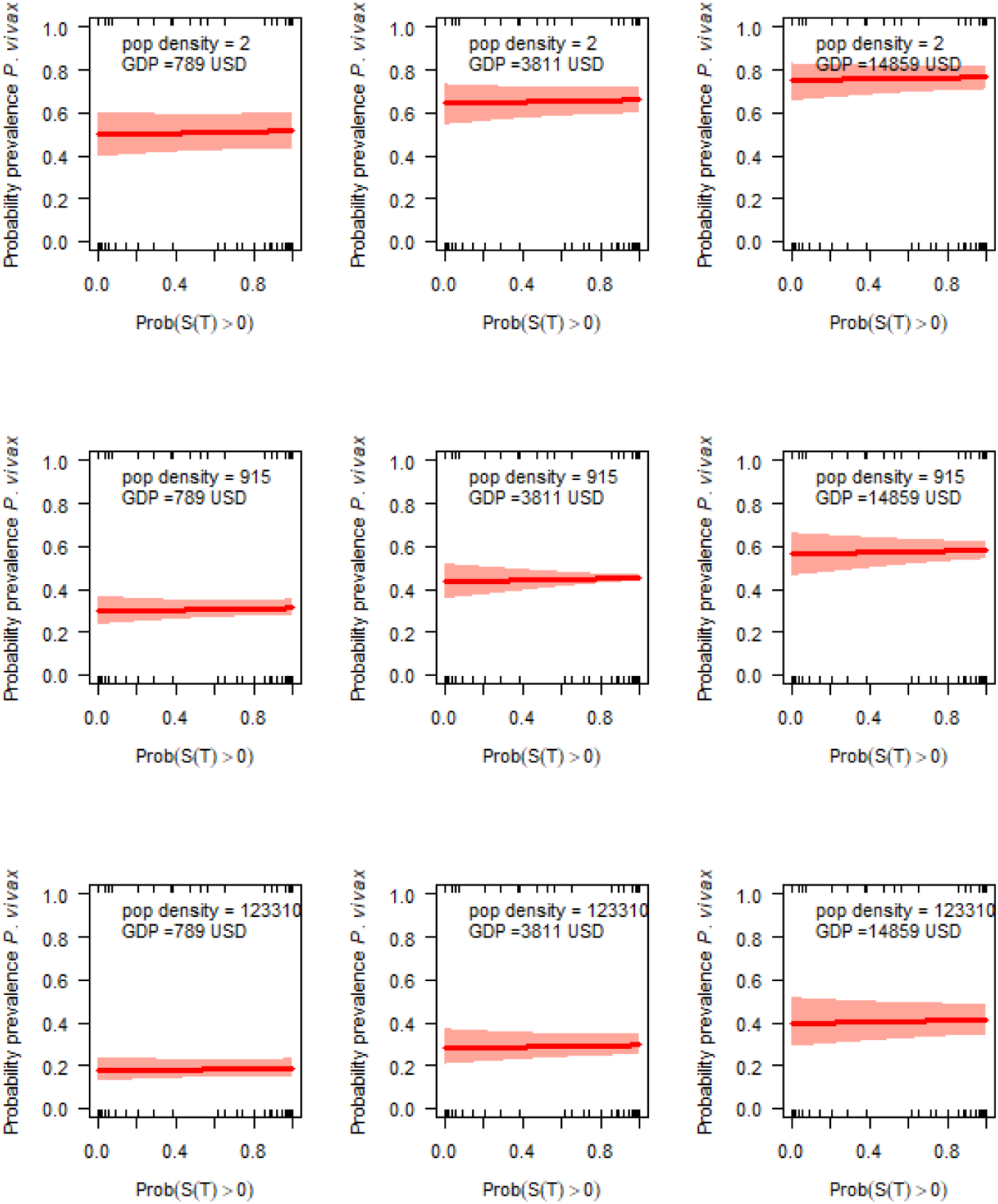
Plots of the probability that *S*(*T*) >0 versus the probability of malaria prevalence by *P. vivax* transmitted by *An. stephensi* in Asia from presence/absence model *Pviv* PA5 (Table 9; SM), for different levels of population density across different rows and different levels of GDP across different columns. Tick-marks are human malaria prevalence data. Lines and shaded areas: mean and 95% CI from GLM.

**Figure B11:**
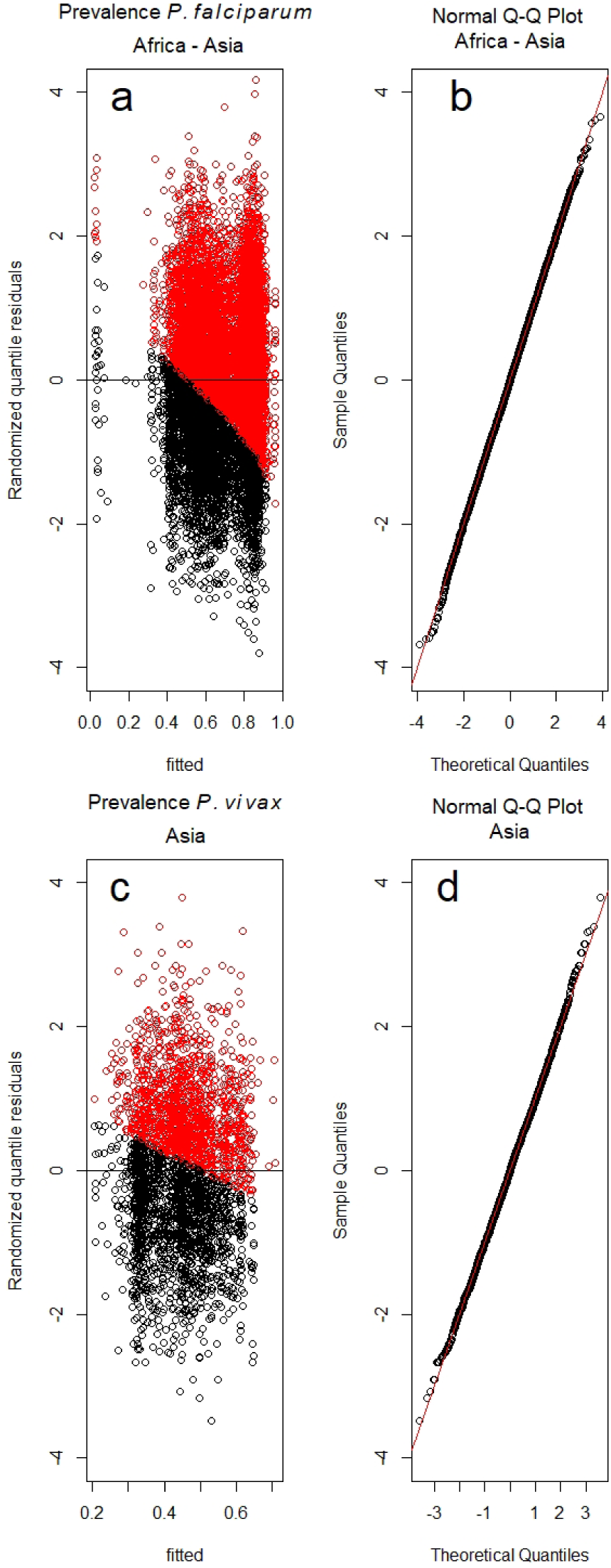
Randomized quantile residuals extracted in R using the *qresid* function in the package *statmod* (Smyth et al., 2019) for the best fitted models (*Pfal* PA5 and *Pviv* PA4) shown in tables 8 and 9 in SM. Randomized Quantile Residuals are interpreted as standards residuals, and should be normally distributed if the assumptions of the underlying model are appropriate for the data. A) Fitted values plotted versus residuals for *P. falciparum* presence (red) and absence (black) prevalence cases. B) Q-Q plot for the quantile residuals for *P. falciparum*. C) fitted values plotted versus residuals for *P. vivax* presence (red) and absence (black) of prevalence cases. D) Q-Q plot for the quantile residuals for *P. vivax*.

### B.6 Summary of the sample posterior distributions, *S*(*T*)

**Figure B12:**
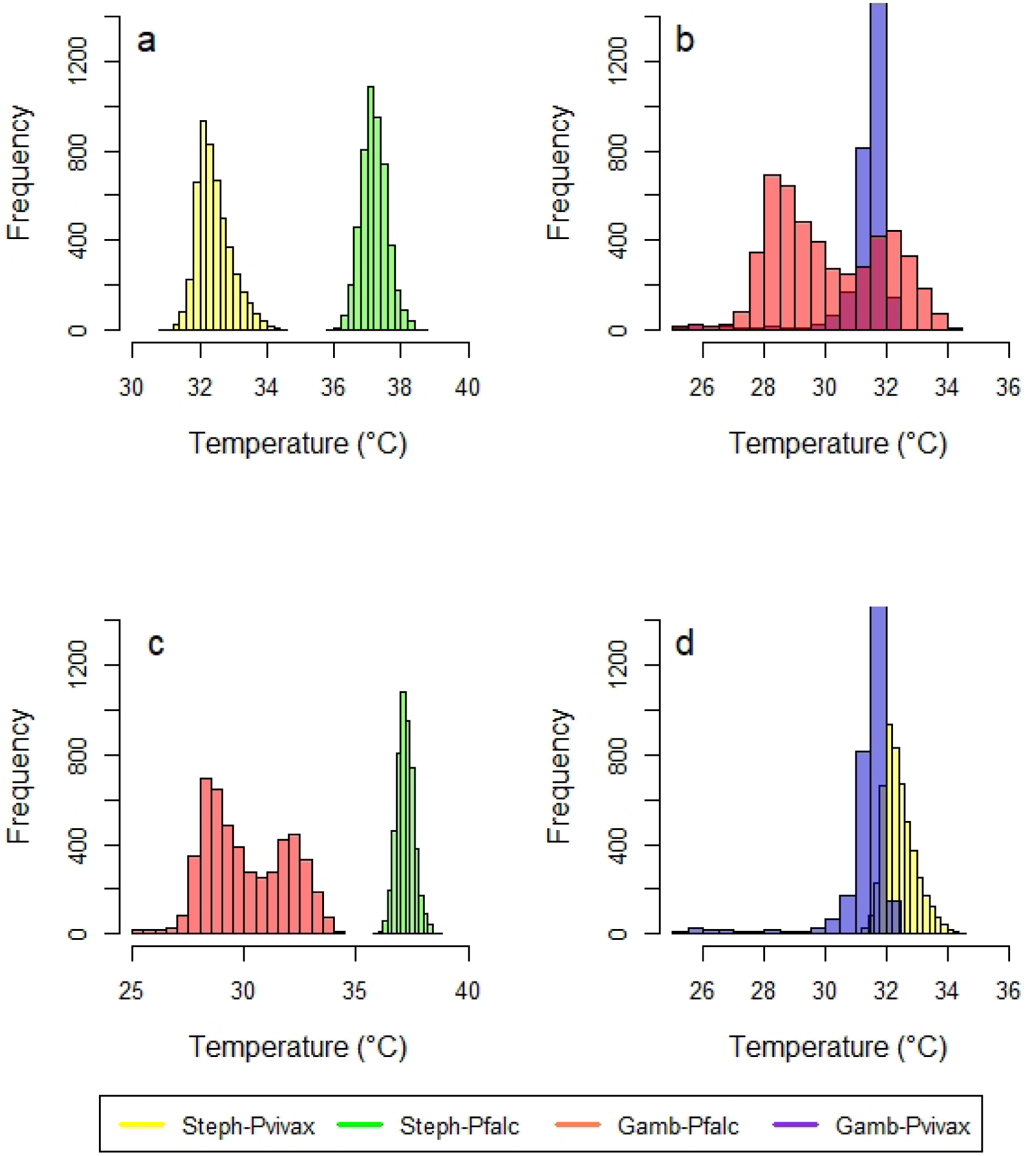
Histograms of the *S*(*T*) posterior distributions at the maximum temperature limit. A) *An. stephensi* with *P. falciparum* versus *An. stephensi* with *P. vivax*. B) *An. gambiae* with *P. falciparum* and *An. gambiae* with *P. vivax*. C) *An. gambiae* with *P. falciparum* versus *An. stephensi* with *P. falciparum*. D) *An. gambiae* with *P. vivax* versus *An. stephensi* with *P. vivax*.

**Figure B13:**
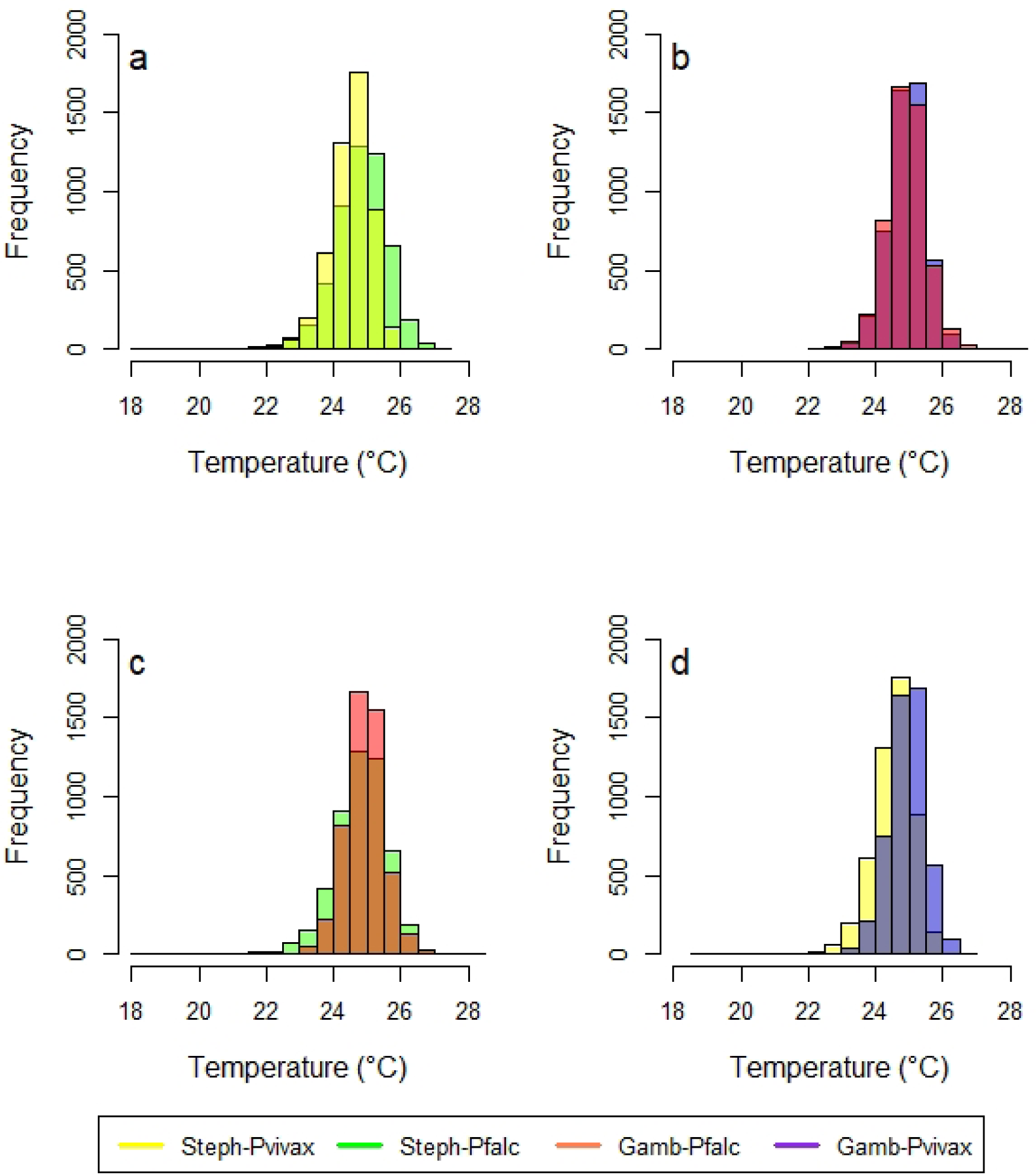
Histograms of the *S*(*T*) posterior distributions at the optimum temperature limit. A) *An. stephensi* with *P. falciparum* versus *An. stephensi* with *P. vivax*. B) *An. gambiae* with *P. falciparum* and *An. gambiae* with *P. vivax*. C) *An. gambiae* with *P. falciparum* versus *An. stephensi* with *P. falciparum*. D) *An. gambiae* with *P. vivax* versus *An. stephensi* with *P. vivax*.

**Figure B14:**
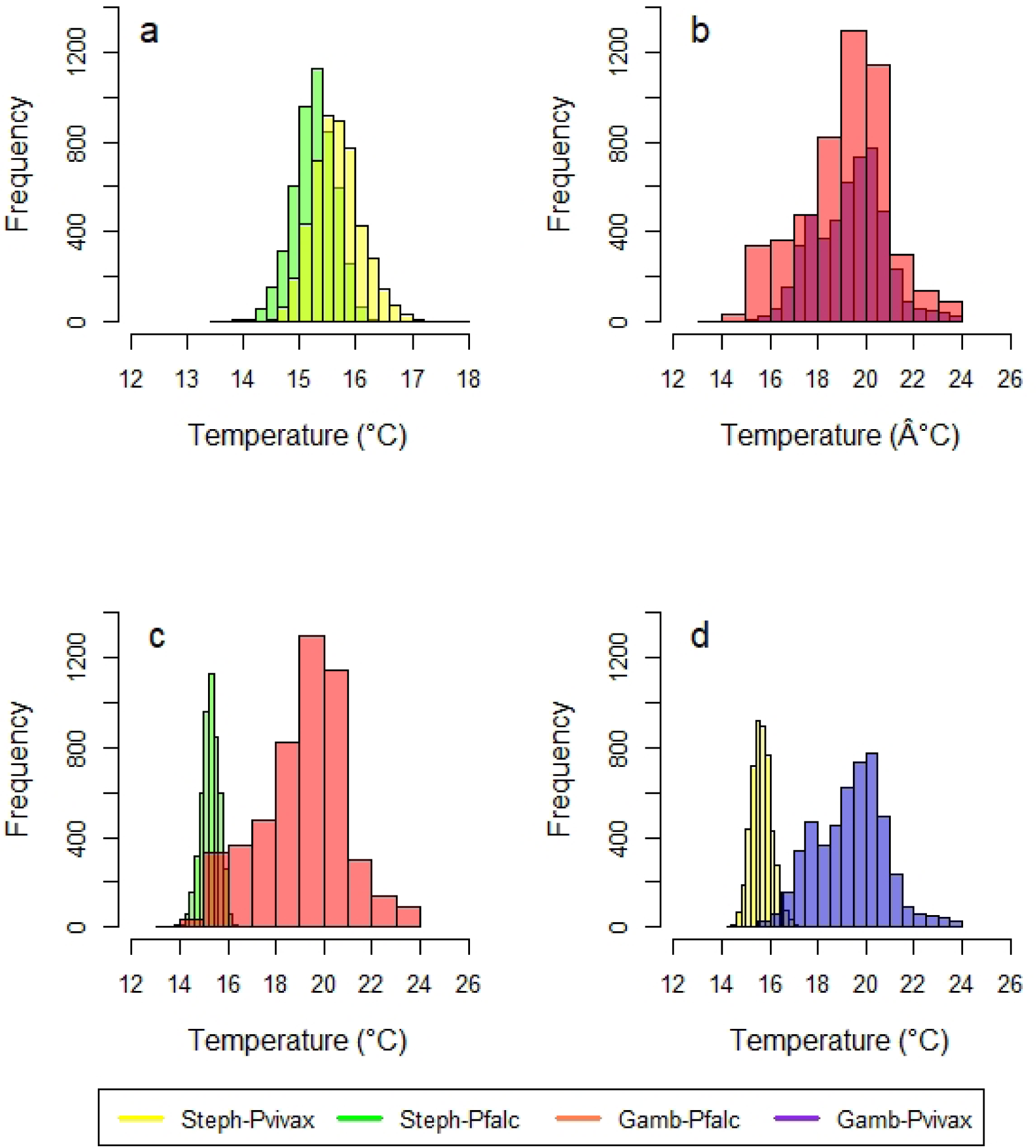
Histograms of the *S*(*T*) posterior distributions at the minimum temperature limit. A) *An. stephensi* with *P. falciparum* versus *An. stephensi* with *P. vivax*. B) *An. gambiae* with *P. falciparum* and *An. gambiae* with *P. vivax*. C) *An. gambiae* with *P. falciparum* versus *An. stephensi* with *P. falciparum*. D) *An. gambiae* with *P. vivax* versus *An. stephensi* with *P. vivax*.

### B.7 Comparison of the mosquito/parasite posterior distributions, *S*(*T*)

**Figure B15:**
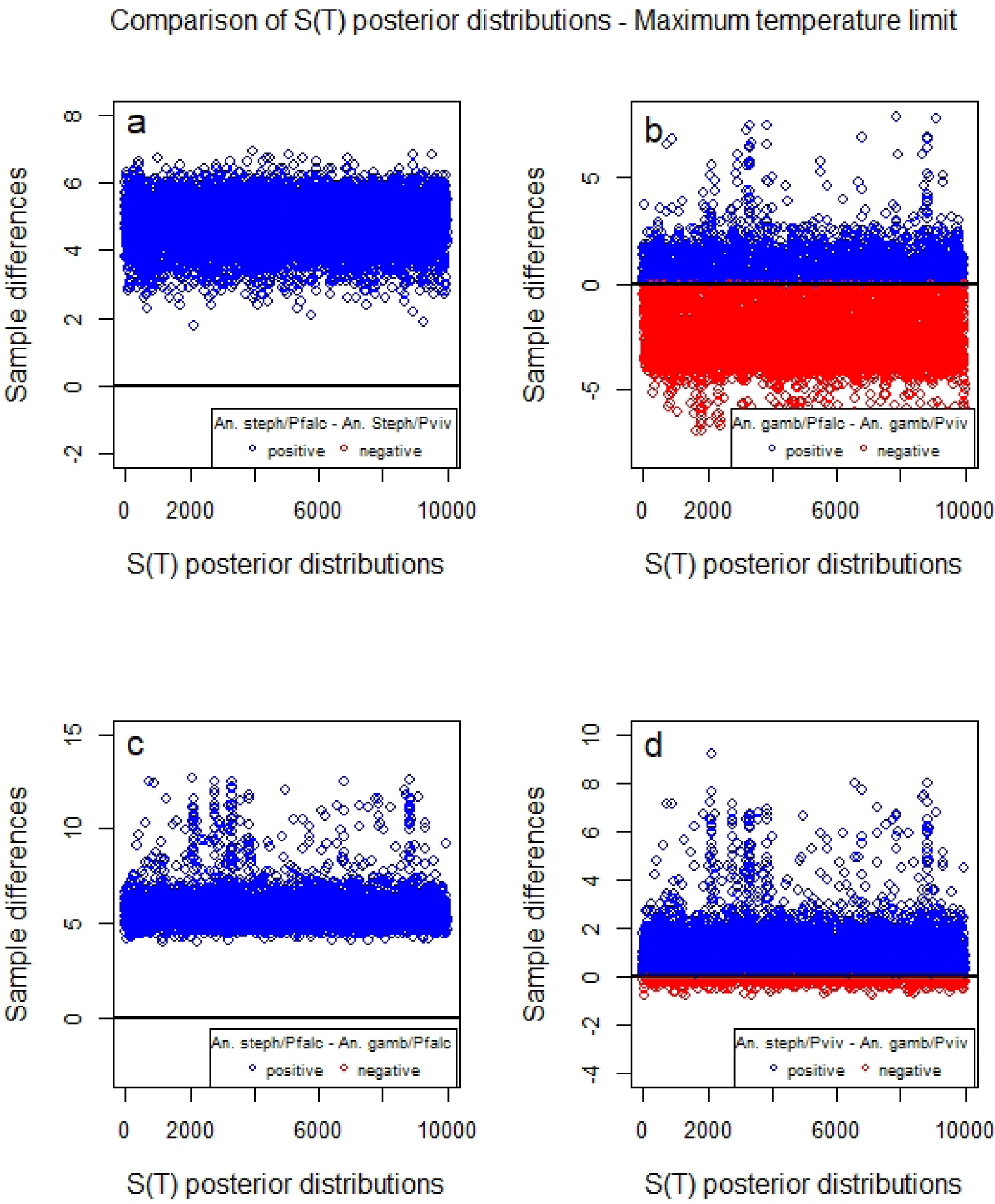
Sample differences of the *S*(*T*) posterior distributions at the maximum temperature limit. A) *An. stephensi* with *P. falciparum* versus *An. stephensi* with *P. vivax* B) *An. gambiae* with *P. falciparum* versus *An. gambiae* with *P. vivax* C) *An. stephensi* with *P. falciparum* versus *An. stephensi* with *P. falciparum*, and D) *An. stephensi* with *P. falciparum* versus *An. gambiae* with *P. vivax*.

**Figure B16:**
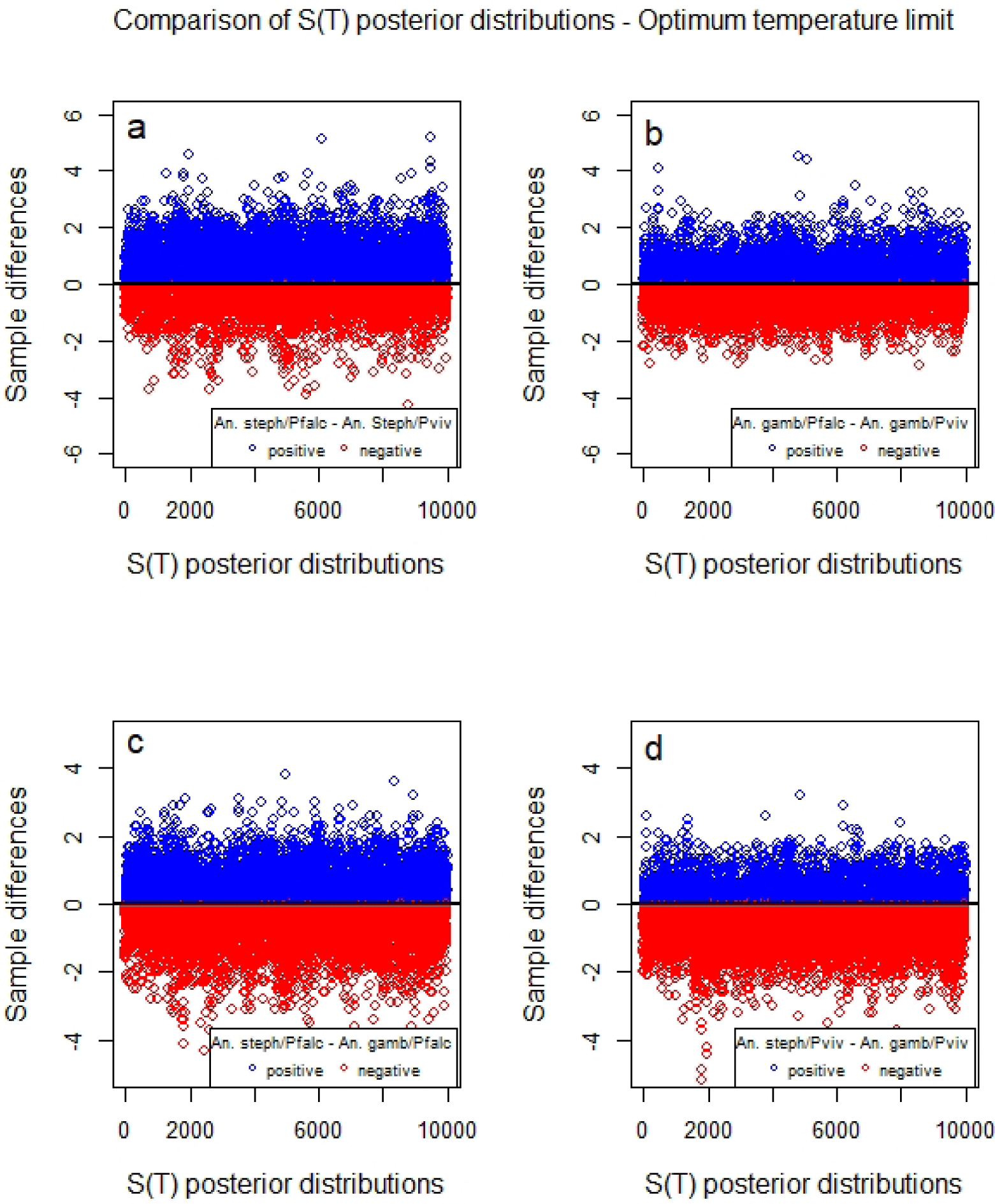
Sample differences of the *S*(*T*) posterior distributions at the optimum temperature limit. A) *An. stephensi* with *P. falciparum* versus *An. stephensi* with *P. vivax* B) *An. gambiae* with *P. falciparum* versus *An. gambiae* with *P. vivax* C) *An. stephensi* with *P. falciparum* versus *An. gambiae* with *P. falciparum*, and D) *An. stephensi* with *P. vivax* versus *An. gambiae* with *P. vivax*.

**Figure B17:**
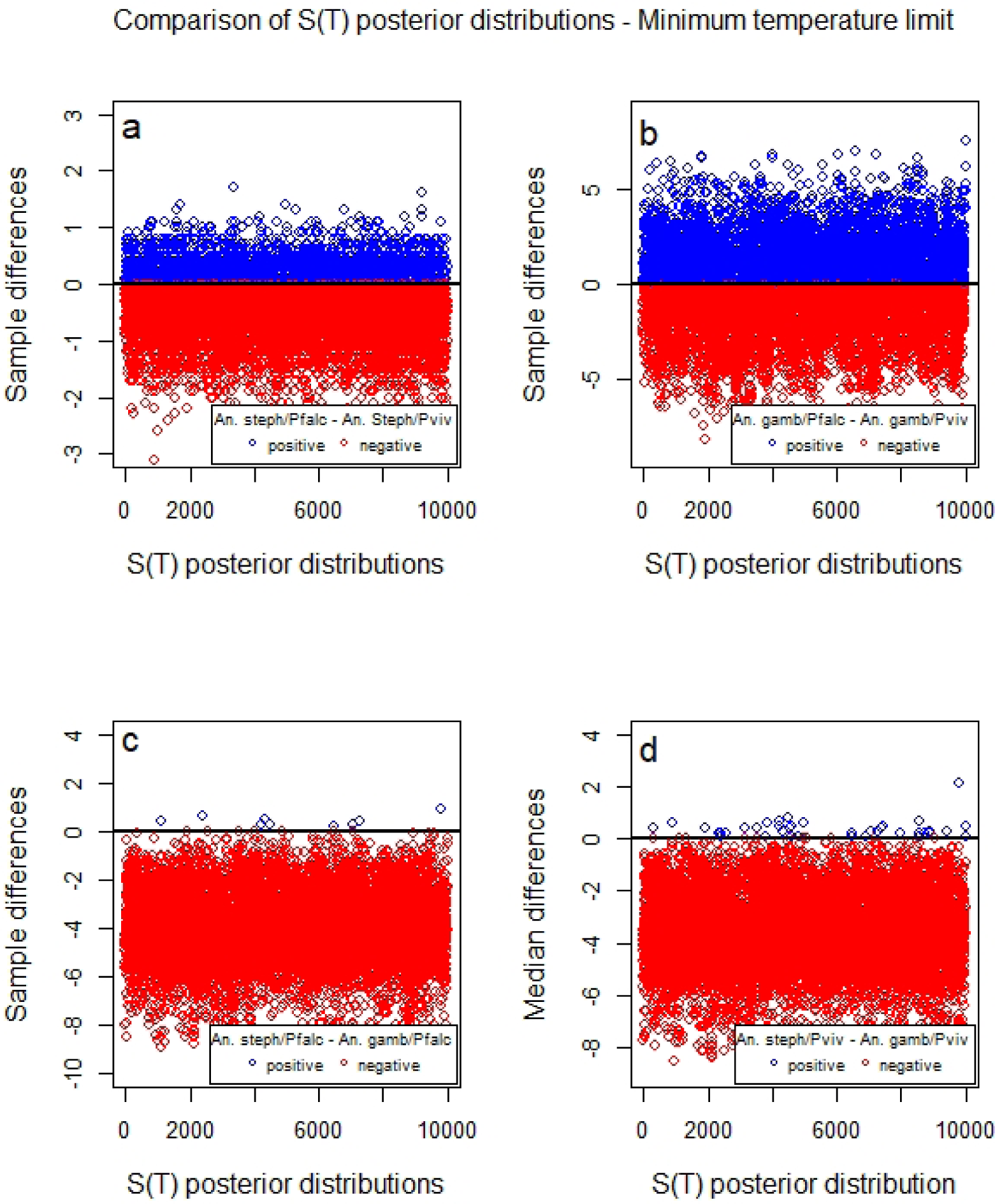
Sample differences of the *S*(*T*) posterior distributions at the minimum temperature limit. A) *An. stephensi* with *P. falciparum* versus *An. stephensi* with *P. vivax* B) *An. gambiae* with *P. falciparum* versus *An. gambiae* with *P. vivax* C) *An. stephensi* with *P. falciparum* versus *An. gambiae* with *P. falciparum*, and D) *An. stephensi* with *P. vivax* versus *An. gambiae* with *P. vivax*.

### B.8 Suitability maps

**Figure B18:**
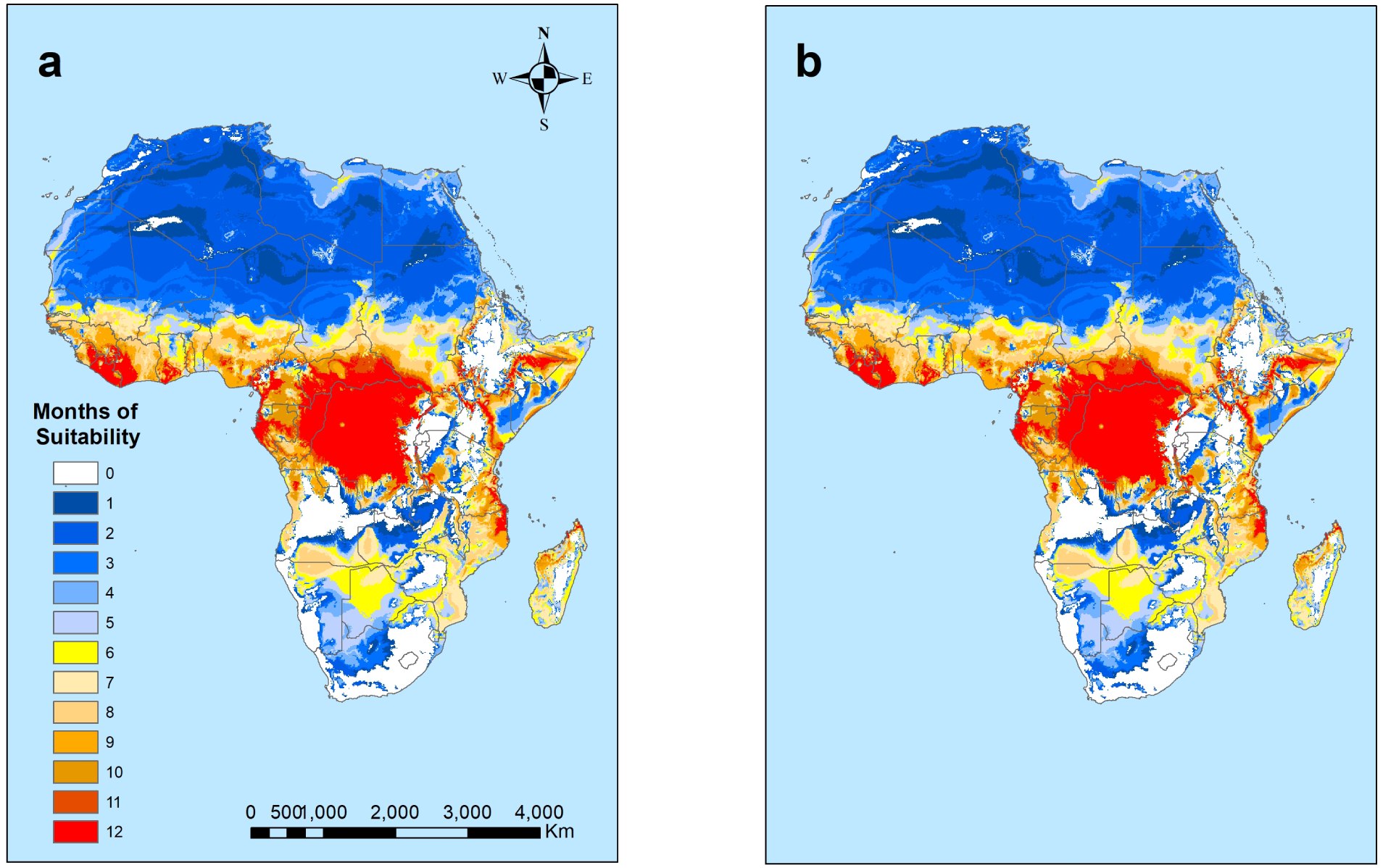
The number of months a year that locations in Africa and Asia are suitable (*S*(*T*)>0) for the transmission of A) *P. falciparum*, B) *P. vivax*, by *An. gambiae* mosquitoes. We define highly suitable temperatures as *S*(*T*)>0.75.

### B.9 Proportion of suitable months for malaria transmission in Africa and Asia

**Figure B19:**
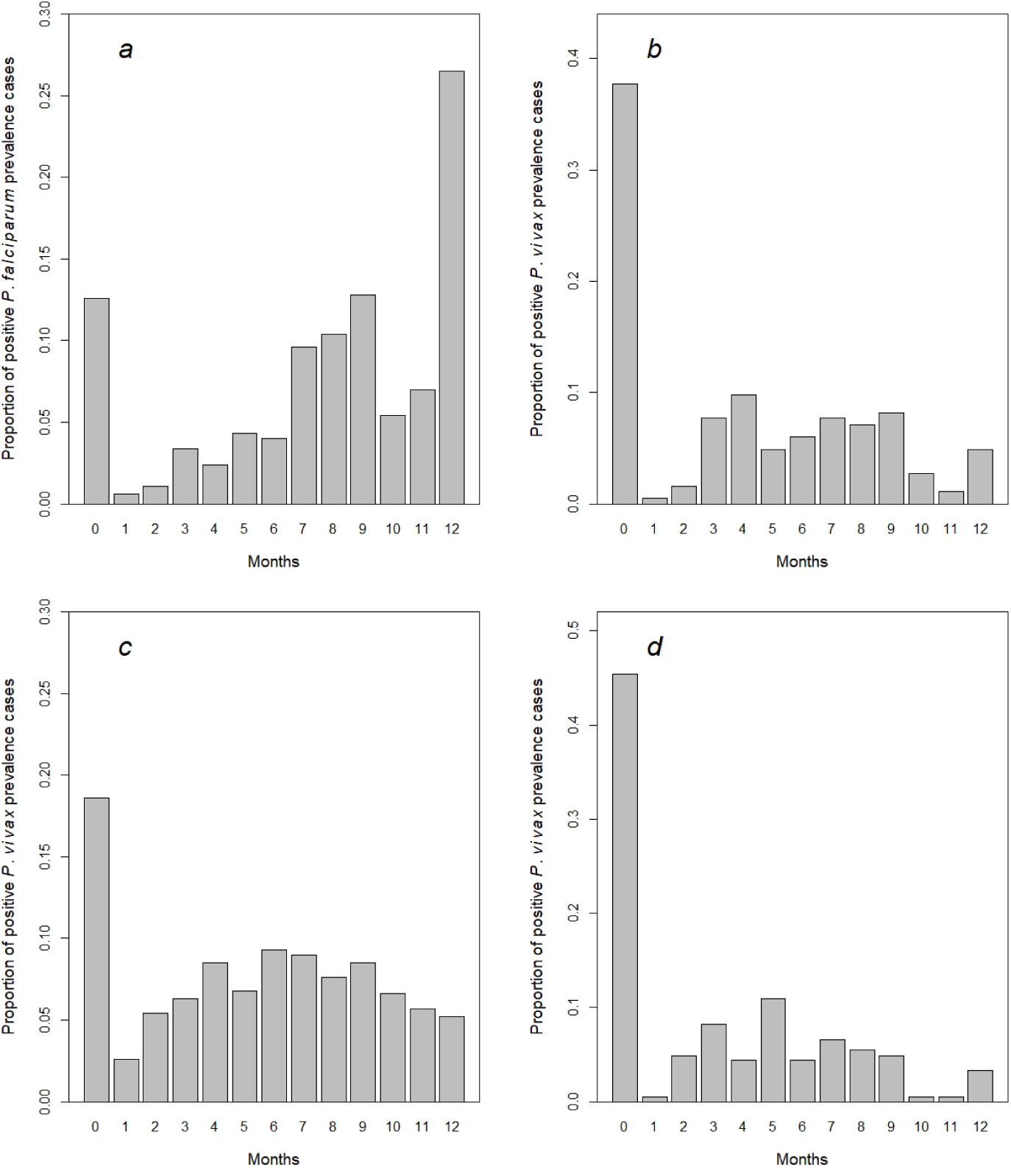
The proportion of number of suitable months for transmission *S*(*T*)>0 in Africa with the presence of A) *P. falciparum* malaria prevalence in area suitable for *An. stephensi* B) *P. vivax* malaria prevalence in area suitable for *An. stephensi* C) *P. falciparum* malaria prevalence in area suitable for *An. gambiae* D) *P. vivax* malaria prevalence in area suitable for *An. gambiae*.

**Figure B20:**
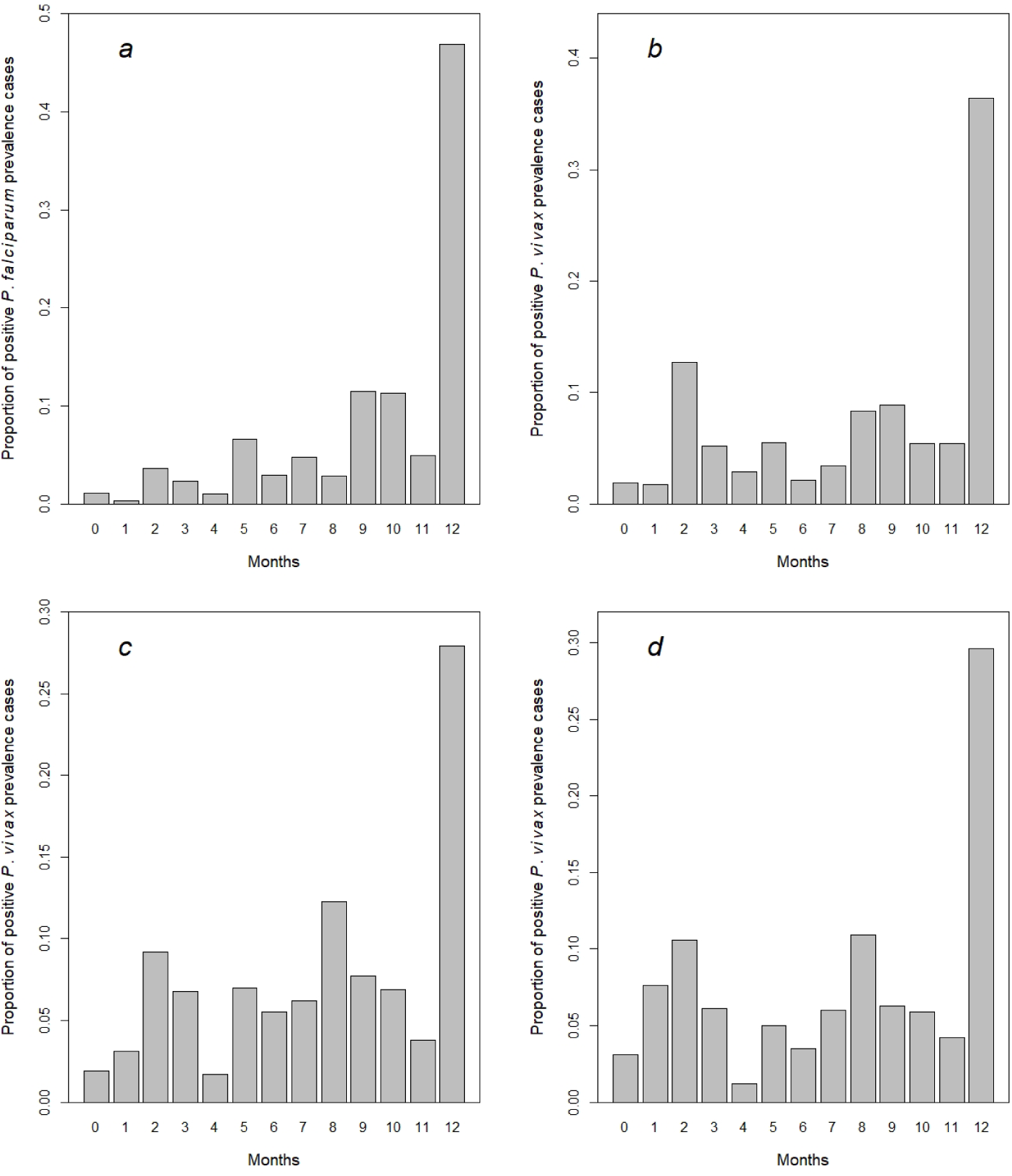
The proportion of the number of suitable months for transmission *S*(*T*)>0 in Asia with the presence of A) *P. falciparum* in area suitable for *An. stephensi* B) *P. vivax* in area suitable for *An. stephensi* C) *P. falciparum* in area suitable for *An. gambiae* D) *P. vivax in area suitable for An. gambiae*.

### B.10 Boxplots of the variables used for model validation for Africa and Asia

**Figure B21:**
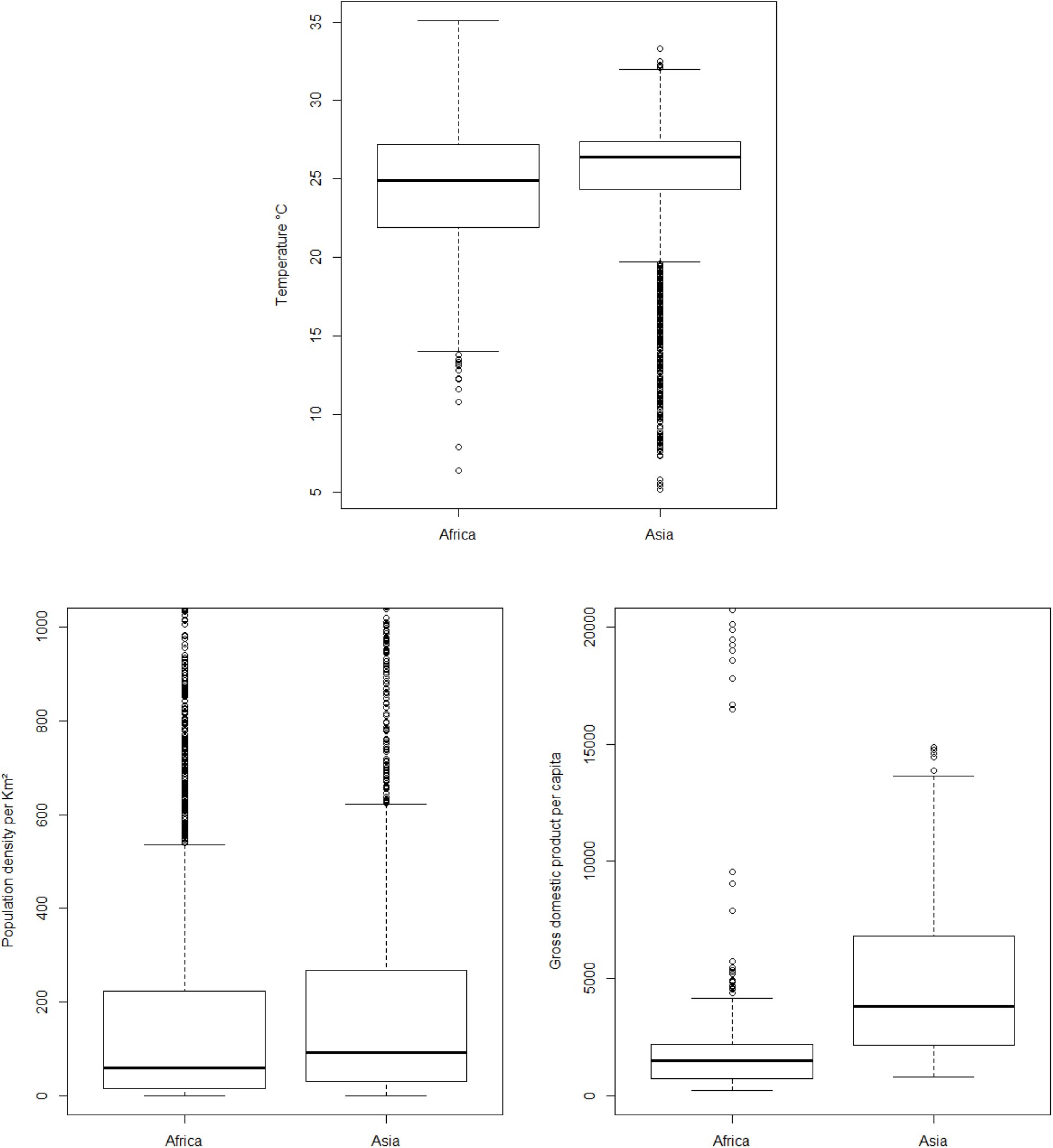
Comparison of the variability of the variables used for model validation between the Africa and Asia regions. A) Temperature expressed in °C B) Population density per km^2^ C) Gross domestic product per capita.

## Terms and abbreviations

BIC: Bayesian information criterion
*D*^2^: Squared Deviance, the proportion of deviance explained by the model
Model pr.: Model probability based on BIC
*η*_*i*_: linear predictor;
*β*_0_: Intercept; *β*_1_,…*β*_*n*_ Regression parameters
lpop den: log(population density)
lGDP: log(per capita gross domestic product)
SGTZ: Probability of *S*(*T*)>0
Pfal: *Plasmodium falciparum*
Pviv: *Plasmodium vivax*
𝟙_*AFR*_: An indicator function that returns 1 if the condition in the subscript is TRUE and zero if FALSE. In our notation, the variable *AFR* is TRUE (=1) if the location is in Africa and if FALSE (=0) otherwise (i.e., if the location is in Asia).

## References

Abram, P. K., Boivin, G., Moiroux, J., and Brodeur, J. (2017). Behavioural effects of temperature on ectothermic animals: unifying thermal physiology and behavioural plasticity. Biological Reviews, 92(4):1859–1876.

Balk, D. L., Deichmann, U., Yetman, G., Pozzi, F., Hay, S. I., and Nelson, A. (2006). Determining global population distribution: methods, applications and data. Advances in parasitology, 62:119–156.

Balkew, M., Mumba, P., Dengela, D., and Yohannes, G. (2019). Geographical distribution of anopheles stephensi in eastern ethiopia. bioRxiv, pages 1–22.

Barik, R. K., Lenka, R. K., Ali, S. M., Gupta, N., Satpathy, A., and Raj, A. (2017). Investigation into the efficacy of geospatial big data visualization tools. In 2017 International Conference on Computing, Communication and Automation (ICCCA), pages 88–93. IEEE.

Bayoh, M. N. (2001). Studies on the development and survival of Anopheles gambiae sensu stricto at various temperatures and relative humidities. PhD thesis, Durham University.

Beck-Johnson, L. M., Nelson, W. A., Paaijmans, K. P., Read, A. F., Thomas, M. B., and Bjørnstad, O. N. (2017). The importance of temperature fluctuations in understanding mosquito population dynamics and malaria risk. Royal Society open science, 4(3):160969.

Burnham, K. P. and Anderson, D. R. (2004). Multimodel Inference: Understanding AIC and BIC in Model Selection. Sociological Methods & Research, 33(2):261–304.

Carter, T. E., Yared, S., Gebresilassie, A., Bonnell, V., Damodaran, L., Lopez, K., Ibrahim, M., Mohammed, S., and Janies, D. (2018). First detection of anopheles stephensi liston, 1901 (diptera: culicidae) in ethiopia using molecular and morphological approaches. Acta tropica, 188:180–186.

Cator, L., Johnson, L. R., Mordecai, E. A., El Moustaid, F., Smallwood, T. R., LaDeau, S. L., Johansson, M. A., Hudson, P. J., Boots, M., Thomas, M. B., et al. (2020). The role of vector trait variation in vector-borne disease dynamics. Frontiers in ecology and evolution.

Changyong, F., Hongyue, W., Naiji, L., Tian, C., Hua, H., Ying, L., et al. (2014). Log-transformation and its implications for data analysis. Shanghai archives of psychiatry, 26(2):105.

Clements, A. N. (2013). The Physiology of Mosquitoes: International Series of Monographs on Pure and Applied Biology: Zoology, volume 17. Elsevier.

Colinet, H., Sinclair, B. J., Vernon, P., and Renault, D. (2015). Insects in fluctuating thermal environments. Annual Review of Entomology, 60:123–140.

Craig, M. H., Snow, R., and le Sueur, D. (1999). A climate-based distribution model of malaria transmission in sub-saharan africa. Parasitology today, 15(3):105–111.

Dell, A. I., Pawar, S., and Savage, V. M. (2011). Systematic variation in the temperature dependence of physiological and ecological traits. Proceedings of the National Academy of Sciences, 108(26):10591–10596.

Dietz, K. (1993). The estimation of the basic reproduction number for infectious diseases. Statistical methods in medical research, 2(1):23–41.

Dunn, P. K. and Smyth, G. K. (1996). Randomized quantile residuals. Journal of Computational and Graphical Statistics, 5(3):236–244.

Dunn, P. K. and Smyth, G. K. (2018). Generalized Linear Models With Examples in R. Springer.

Fang, J. and McCutchan, T. F. (2002). Malaria: Thermoregulation in a parasite’s life cycle. Nature, 418(6899):742.

Faulde, M. K., Rueda, L. M., and Khaireh, B. A. (2014). First record of the asian malaria vector anopheles stephensi and its possible role in the resurgence of malaria in djibouti, horn of africa. Acta tropica, 139:39–43.

Fick, S. E. and Hijmans, R. J. (2017). Worldclim 2: new 1-km spatial resolution climate surfaces for global land areas. International journal of climatology, 37(12):4302–4315.

Geissbühler, Y., Chaki, P., Emidi, B., Govella, N. J., Shirima, R., Mayagaya, V., Mtasiwa, D., Mshinda, H., Fillinger, U., Lindsay, S. W., et al. (2007). Interdependence of domestic malaria prevention measures and mosquito-human interactions in urban dar es salaam, tanzania. Malaria journal, 6(1):126.

Giner, G. and Smyth, G. K. (2016). statmod: probability calculations for the inverse gaussian distribution. R Journal, 8(1):339–351.

Hijmans, R. J., Van Etten, J., Cheng, J., Mattiuzzi, M., Sumner, M., Greenberg, J. A., Lamigueiro, O. P., Bevan, A., Racine, E. B., Shortridge, A., et al. (2015). Package ‘raster’. R package.

Holme, P. and Masuda, N. (2015). The basic reproduction number as a predictor for epidemic outbreaks in temporal networks. PloS one, 10(3):e0120567.

James, S. L., Gubbins, P., Murray, C. J., and Gakidou, E. (2012). Developing a comprehensive time series of gdp per capita for 210 countries from 1950 to 2015. Population health metrics, 10(1):12.

Johnson, L. R., Ben-Horin, T., Lafferty, K. D., McNally, A., Mordecai, E., Paaijmans, K. P., Pawar, S., and Ryan, S. J. (2015). Understanding uncertainty in temperature effects on vector-borne disease: a bayesian approach. Ecology, 96(1):203–213.

Kern, P., Cramp, R. L., and Franklin, C. E. (2015). Physiological responses of ectotherms to daily temperature variation. Journal of Experimental Biology, 218(19):3068–3076.

Kirby, M. and Lindsay, S. (2004). Responses of adult mosquitoes of two sibling species, anopheles arabiensis and a. gambiae ss (diptera: Culicidae), to high temperatures. Bulletin of Entomological Research, 94(5):441–448.

Lahondére, C. and Lazzari, C. R. (2012). Mosquitoes cool down during blood feeding to avoid overheating. Current biology, 22(1):40–45.

Lunde, T. M., Bayoh, M. N., and Lindtjørn, B. (2013). How malaria models relate temperature to malaria transmission. Parasites & vectors, 6(1):20.

Lyons, C. L., Coetzee, M., Terblanche, J. S., and Chown, S. L. (2012). Thermal limits of wild and laboratory strains of two african malaria vector species, anopheles arabiensis and anopheles funestus. Malaria Journal, 11(1):226.

Mahmood, F. (1997). Life-table attributes of anopheles albimanus (wiedemann) under controlled laboratory conditions. Journal of vector ecology: journal of the Society for Vector Ecology, 22(2):103–108.

Miazgowicz, K., Shocket, M., Ryan, S., Villena, O., Hall, R., Owen, J., Adanlawo, T., Balaji, K., Johson, L., Mordecai, E., and Murdock, C. (2020). Age influences the thermal suitability of plasmodium falciparum transmission in the asian malaria vector anopheles stephensi. Procedings of the Royal Society B, RSPB-2020-1093.R1.

Mordecai, E. A., Caldwell, J. M., Grossman, M. K., Lippi, C. A., Johnson, L. R., Neira, M., Rohr, J. R., Ryan, S. J., Savage, V., Shocket, M. S., et al. (2019). Thermal biology of mosquito-borne disease. Ecology letters.

Mordecai, E. A., Cohen, J. M., Evans, M. V., Gudapati, P., Johnson, L. R., Lippi, C. A., Miazgowicz, K., Murdock, C. C., Rohr, J. R., Ryan, S. J., et al. (2017). Detecting the impact of temperature on transmission of zika, dengue, and chikungunya using mechanistic models. PLoS neglected tropical diseases, 11(4):e0005568.

Mordecai, E. A., Paaijmans, K. P., Johnson, L. R., Balzer, C., Ben-Horin, T., de Moor, E., McNally, A., Pawar, S., Ryan, S. J., Smith, T. C., et al. (2013). Optimal temperature for malaria transmission is dramatically lower than previously predicted. Ecology letters, 16(1):22–30.

Paaijmans, K. P. and Thomas, M. B. (2011). The influence of mosquito resting behaviour and associated microclimate for malaria risk. Malaria Journal, 10(1):183.

Parham, P. E. and Michael, E. (2009). Modeling the effects of weather and climate change on malaria transmission. Environmental health perspectives, 118(5):620–626.

Pfeffer, D. A., Lucas, T. C., May, D., Harris, J., Rozier, J., Twohig, K. A., Dalrymple, U., Guerra, C. A., Moyes, C. L., Thorn, M., et al. (2018). malariaatlas: an r interface to global malariometric data hosted by the malaria atlas project. Malaria journal, 17(1):1–10.

Plummer, M. (2016). rjags: Bayesian Graphical Models using MCMC. R package version 4–6.

R Development Core Team (2017). R: A Language and Environment for Statistical Computing. R Foundation for Statistical Computing, Vienna, Austria.

Scott, L. M. and Janikas, M. V. (2010). Spatial statistics in arcgis. In Handbook of applied spatial analysis, pages 27–41. Springer.

Shapiro, L. L., Whitehead, S. A., and Thomas, M. B. (2017). Quantifying the effects of temperature on mosquito and parasite traits that determine the transmission potential of human malaria. PLoS biology, 15(10):e2003489.

Sinclair, B. J., Marshall, K. E., Sewell, M. A., Levesque, D. L., Willett, C. S., Slotsbo, S., Dong, Y., Harley, C. D., Marshall, D. J., Helmuth, B. S., et al. (2016). Can we predict ectotherm responses to climate change using thermal performance curves and body temperatures? Ecology Letters, 19(11):1372–1385.

Sinka, M. E., Bangs, M. J., Manguin, S., Chareonviriyaphap, T., Patil, A. P., Temperley, W. H., Gething, P. W., Elyazar, I. R., Kabaria, C. W., Harbach, R. E., et al. (2011). The dominant anopheles vectors of human malaria in the asia-pacific region: occurrence data, distribution maps and bionomic précis. Parasites & vectors, 4(1):89.

Sinka, M. E., Bangs, M. J., Manguin, S., Coetzee, M., Mbogo, C. M., Hemingway, J., Patil, A. P., Temperley, W. H., Gething, P. W., Kabaria, C. W., et al. (2010). The dominant anopheles vectors of human malaria in africa, europe and the middle east: occurrence data, distribution maps and bionomic précis. Parasites & vectors, 3(1):117.

Sinka, M. E., Bangs, M. J., Manguin, S., Rubio-Palis, Y., Chareonviriyaphap, T., Coetzee, M., Mbogo, C. M., Hemingway, J., Patil, A. P., Temperley, W. H., et al. (2012). A global map of dominant malaria vectors. Parasites & vectors, 5(1):69.

Snow, R. W., Guerra, C. A., Noor, A. M., Myint, H. Y., and Hay, S. I. (2005). The global distribution of clinical episodes of plasmodium falciparum malaria. Nature, 434(7030):214.

Takken, W. and Lindsay, S. (2019). Increased threat of urban malaria from anopheles stephensi mosquitoes, africa. Emerging infectious diseases, 25(7):1431.

Taylor, R. A., Ryan, S. J., Lippi, C. A., Hall, D. G., Narouei-Khandan, H. A., Rohr, J. R., and Johnson, L. R. (2019). Predicting the fundamental thermal niche of crop pests and diseases in a changing world: a case study on citrus greening. Journal of Applied Ecology, 56(8):2057–2068.

Tesla, B., Demakovsky, L. R., Mordecai, E. A., Ryan, S. J., Bonds, M. H., Ngonghala, C. N., Brindley, M. A., and Murdock, C. C. (2018). Temperature drives zika virus transmission: evidence from empirical and mathematical models. Proceedings of the Royal Society B: Biological Sciences, 285(1884):20180795.

Tizifa, T. A., Kabaghe, A. N., McCann, R. S., van den Berg, H., Van Vugt, M., and Phiri, K. S. (2018). Prevention efforts for malaria. Current tropical medicine reports, 5(1):41–50.

World Health Organization (2008). World malaria report 2008. Technical report, WHO Geneva.

World Health Organization (2018). World malaria report 2018. Technical report, WHO Geneva.

World Health Organization (2019). Vector alert: Anopheles stephensi invasion and spread: Horn of africa, the republic of the sudan and surrounding geographical areas, and sri lanka: information note. Technical report, World Health Organization.

## References

Afrane, Y. A., Little, T. J., Lawson, B. W., Githeko, A. K., and Yan, G. (2008). Deforestation and vectorial capacity of anopheles gambiae giles mosquitoes in malaria transmission, kenya. Emerging Infectious Diseases, 14(10):1533.

Afrane, Y. A., Zhou, G., Lawson, B. W., Githeko, A. K., and Yan, G. (2007). Life-table analysis of anopheles arabiensis in western kenya highlands: effects of land covers on larval and adult survivorship. The American journal of tropical medicine and hygiene, 77(4):660–666.

Barreaux, A. M., Barreaux, P., Thievent, K., and Koella, J. (2016). Larval environment influences vector competence of the malaria mosquito anopheles gambiae. Malaria World J, 7(8):1–6.

Bayoh, M. and Lindsay, S. (2003). Effect of temperature on the development of the aquatic stages of anopheles gambiae sensu stricto (diptera: Culicidae). Bulletin of entomological research, 93(5):375–381.

Boyd, M. F. et al. (1932). Studies on plasmodium vivax. 2. the influence of temperature on the duration of the extrinsic incubation period. American Journal of Hygiene, 16(3).

Boyd, M. F. and Stratman-Thomas, W. K. (1933). A note on the transmission of quartan malaria by anopheles quadrimaculatus1. The American Journal of Tropical Medicine and Hygiene, 1(3):265–271.

Carter, T. E., Yared, S., Gebresilassie, A., Bonnell, V., Damodaran, L., Lopez, K., Ibrahim, M., Mohammed, S., and Janies, D. (2018). First detection of anopheles stephensi liston, 1901 (diptera: culicidae) in ethiopia using molecular and morphological approaches. Acta tropica, 188:180–186.

Christiansen-Jucht, C. D., Parham, P. E., Saddler, A., Koella, J. C., and Basáñez, M.-G. (2015). Larval and adult environmental temperatures influence the adult reproductive traits of anopheles gambiae ss. Parasites & vectors, 8(1):456.

Eling, W., Hooghof, J., van de Vegte-Bolmer, M., Sauerwein, R., and Van Gemert, G. (2001). Tropical temperatures can inhibit development of the human malaria parasite plasmodium falci-parum in the mosquito. In Proceedings of the Section Experimental and Applied Entomology-Netherlands Entomological Society, volume 12, pages 151–156.

Grieco, J. P., Achee, N. L., Briceno, I., King, R., Andre, R., Roberts, D., and Rejmankova, E. (2003). Comparison of life table attributes from newly established colonies of anopheles albimanus and anopheles vestitipennis in northern belize. Journal of vector ecology, 28:200–207.

Kirby, M. J. and Lindsay, S. W. (2009). Effect of temperature and inter-specific competition on the development and survival of anopheles gambiae sensu stricto and an. arabiensis larvae. Acta tropica, 109(2):118–123.

Knowles, R., Basu, B., et al. (1943). Laboratory studies on the infectivity of anopheles stephensi. Journal of the Malaria Institute of India, 5(1).

Lardeux, F., Loayza, P., Bouchité, B., and Chavez, T. (2007). Host choice and human blood index of anopheles pseudopunctipennis in a village of the andean valleys of bolivia. Malaria journal, 6(1):8.

Lardeux, F. J., Tejerina, R. H., Quispe, V., and Chavez, T. K. (2008). A physiological time analysis of the duration of the gonotrophic cycle of anopheles pseudopunctipennis and its implications for malaria transmission in bolivia. Malaria journal, 7(1):141.

Love, G. and Whelchel, J. (1957). Lethal effects of high temperatures on the immature stages of anopheles quadrimaculatus. Ecology, 38(4):570–576.

Lyons, C. L., Coetzee, M., and Chown, S. L. (2013). Stable and fluctuating temperature effects on the development rate and survival of two malaria vectors, anopheles arabiensis and anopheles funestus. Parasites & vectors, 6(1):104.

Mahande, A., Mosha, F., Mahande, J., and Kweka, E. (2007). Feeding and resting behaviour of malaria vector, anopheles arabiensis with reference to zooprophylaxis. Malaria journal, 6(1):100.

Maharaj, R. (1996). Effects of Temperature on Members of the Anopheles Gambiae Complex (Diptera: Culicidae) in South Africa: Implications for Malaria Transmission and Control. PhD thesis, University of Natal.

Murdock, C., Sternberg, E., and Thomas, M. (2016). Malaria transmission potential could be reduced with current and future climate change. Scientific reports, 6:27771.

Neafsey, D. E., Waterhouse, R. M., Abai, M. R., Aganezov, S. S., Alekseyev, M. A., Allen, J. E., Amon, J., Arca, B., Arensburger, P., Artemov, G., et al. (2015). Highly evolvable malaria vectors: the genomes of 16 anopheles mosquitoes. Science, 347(6217):1258522.

Olayemi, I. K. and Ande, A. T. (2009). Life table analysis of anopheles gambiae (diptera: Culici-dae) in relation to malaria transmission. Journal of Vector Borne Diseases, 46(4):295–299.

O’Loughlin, S. M., Magesa, S., Mbogo, C., Mosha, F., Midega, J., Lomas, S., and Burt, A. (2014). Genomic analyses of three malaria vectors reveals extensive shared polymorphism but contrasting population histories. Molecular biology and evolution, 31(4):889–902.

Paaijmans, K. P., Blanford, S., Chan, B. H., and Thomas, M. B. (2011). Warmer temperatures reduce the vectorial capacity of malaria mosquitoes. Biology letters, 8(3):465–468.

Paaijmans, K. P., Cator, L. J., and Thomas, M. B. (2013a). Temperature-dependent pre-bloodmeal period and temperature-driven asynchrony between parasite development and mosquito biting rate reduce malaria transmission intensity. PloS one, 8(1):e55777.

Paaijmans, K. P., Heinig, R. L., Seliga, R. A., Blanford, J. I., Blanford, S., Murdock, C. C., and Thomas, M. B. (2013b). Temperature variation makes ectotherms more sensitive to climate change. Global change biology, 19(8):2373–2380.

Ponçon, N., Toty, C., L’Ambert, G., Le Goff, G., Brengues, C., Schaffner, F., and Fontenille, D. (2007). Biology and dynamics of potential malaria vectors in southern france. Malaria Journal, 6(1):18.

Rios, L. M. and Connelly, C. R. (2012). Common malaria mosquito anopheles quadrimaculatus say (insecta: Diptera: Culicidea). University of Florida. EENY-419.

Sharma, V. and Dev, V. (2015). Biology & control of anopheles culicifacies giles 1901. The Indian journal of medical research, 141(5):525.

Shute, P. and Maryon, M. (1952). A study of human malaria oocysts as an aid to species diagnosis. Transactions of the Royal Society of Tropical Medicine and Hygiene, 46(3):275–292.

Siddons, L. et al. (1944). The experimental transmission of quartan malaria by anopheles culicifacies giles. Journal of the Malaria Institute of India, 5(3):361–73.

Smyth, G., Hu, Y., Dunn, P., Phipson, B., Chen, Y., and Smyth, M. G. (2019). Package ‘statmod’.

Stratman-Thomas, W. K. (1940). The influence of temperature on plasmodium vivax. The American Journal of Tropical Medicine and Hygiene, 1(5):703–715.

Thomas, S., Ravishankaran, S., Justin, N. J. A., Asokan, A., Kalsingh, T. M. J., Mathai, M. T., Valecha, N., Montgomery, J., Thomas, M. B., and Eapen, A. (2018). Microclimate variables of the ambient environment deliver the actual estimates of the extrinsic incubation period of plasmodium vivax and plasmodium falciparum: a study from a malaria-endemic urban setting, chennai in india. Malaria journal, 17(1):201.

Vélez, I. D. (2008). Desarrollo de un sistema de alerta temprana para la malaria en colombia.

